## Supplementary File 1 for "*Cortex cis*-regulatory switches establish scale colour identity and pattern diversity in *Heliconius*"

**Supplementary File 1**: *Domeless* duplication history in the Lepidoptera.

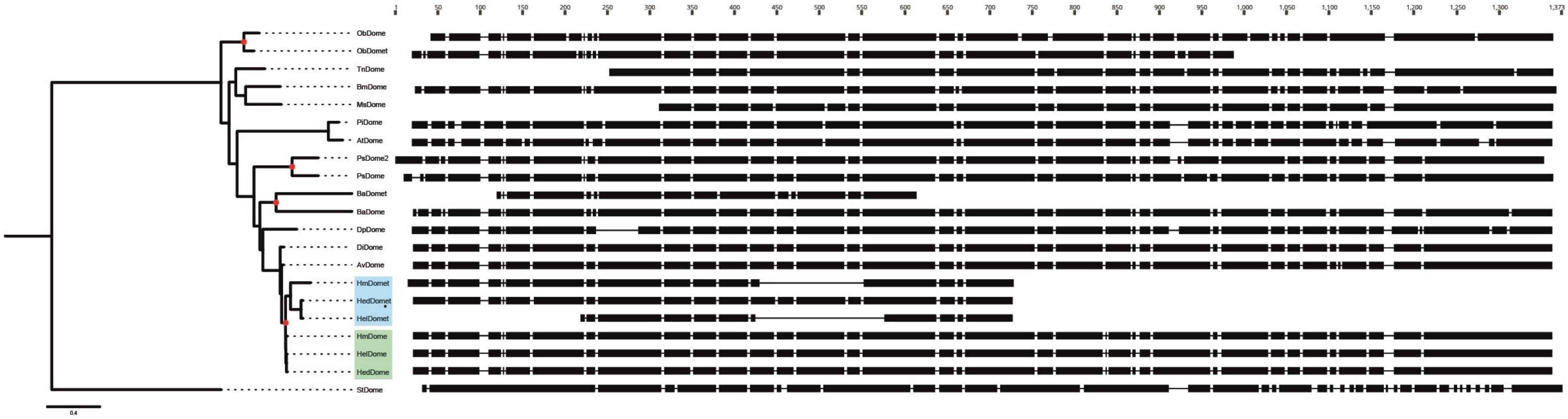


**Figure S1**: Maximum likelihood tree based on lepidopteran *dome* amino acid sequences. The *Heliconius* duplications are highlighted in blue (*domet*) and green (*dome*). Putative duplication events are shown with a red circle. Protein alignments are shown alongside each species, illustrating several C-terminal truncation events in the duplicated sequences. *H. melpomene* (Hm), *H.erato demophoon* (Hed), *Operophtera brumata* (Ob), *Trichoplusia ni* (Tn), *Bombyx mori* (Bm), *Manduca sexta* (Ms), *Plodia interpunctella* (Pi), *Amyeolis transitella* (At), *Phoebis sennae* (Ps), *Bicyclus anynana* (Ba), *Danaus plexippus* (Dp), *Dryas iulia* (Di), *Agraulis vanillae* (Av), *Heliconius erato lativitta* (Hel). As a trichopteran outgroup we used a recently published Pacbio assembly of *Stenopsyche tienmushanensis* (St) (Luo et al., 2018)
