## Supplementary File 2 for "*Cortex cis*-regulatory switches establish scale colour identity and pattern diversity in *Heliconius*"

**Supplementary File 2**: Bi-cistronic transcription of *domeless* and *washout* is conserved across the lepidoptera.

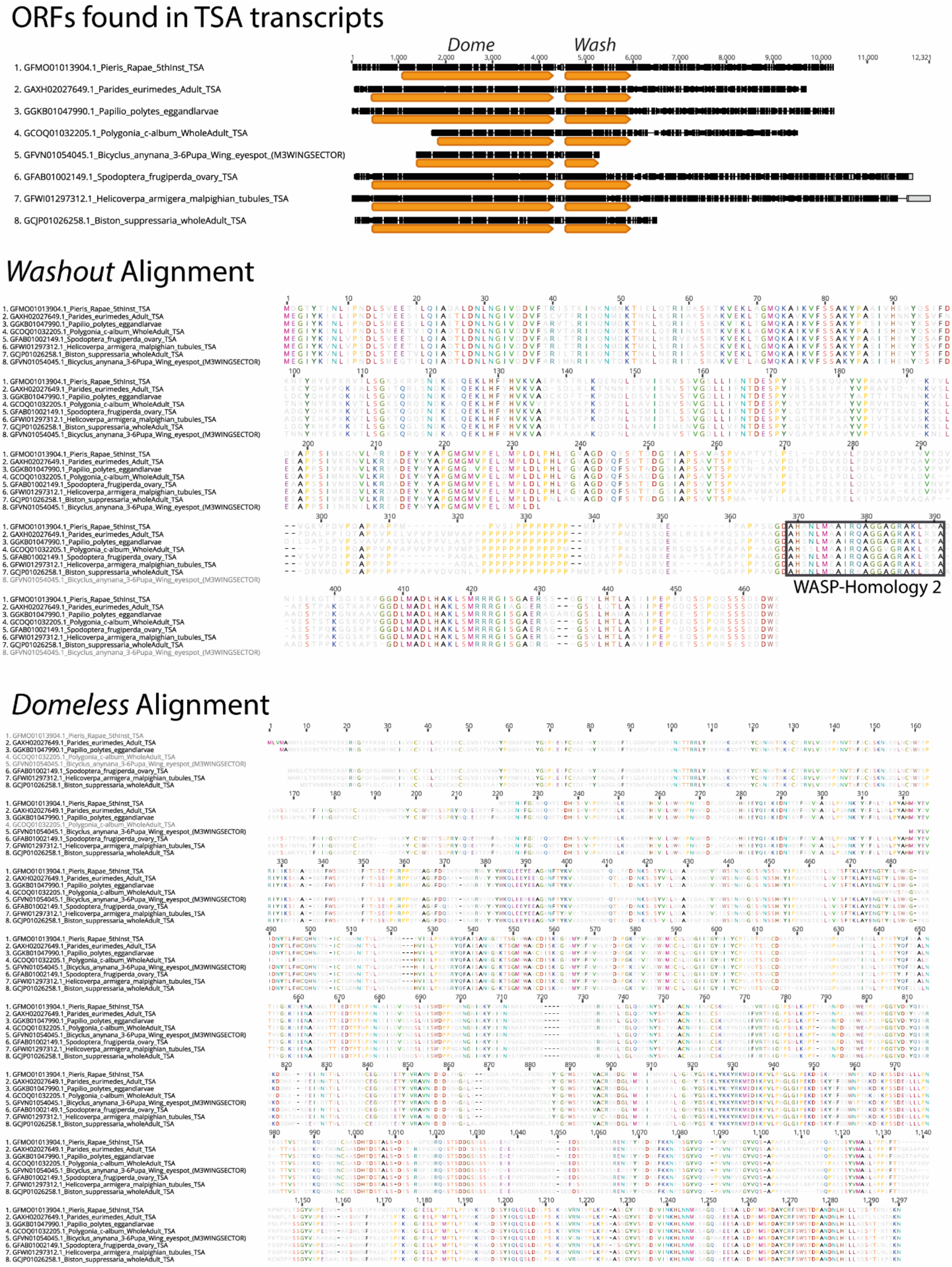


**Figure S2.1:** To examine the conservation of *dome/wash* bi-cistronic transcription in the Lepidoptera we performed BLASTn searches using the previously annotated *dome* transcripts from the *H. melpomene* genome (Hmel2) found on Lepbase (Challi et al., 2016), against the Transcription Shotgun Assembly (TSA) sequence archive on NCBI. We recovered several assembled transcripts containing both the *dome* and *wash* ORFs in various divergent lepidoptera. The positions of *dome* and *wash* ORFs are shown (arrows blocks in TSA transcript) as well as the encoded Wash and Dome proteins below.

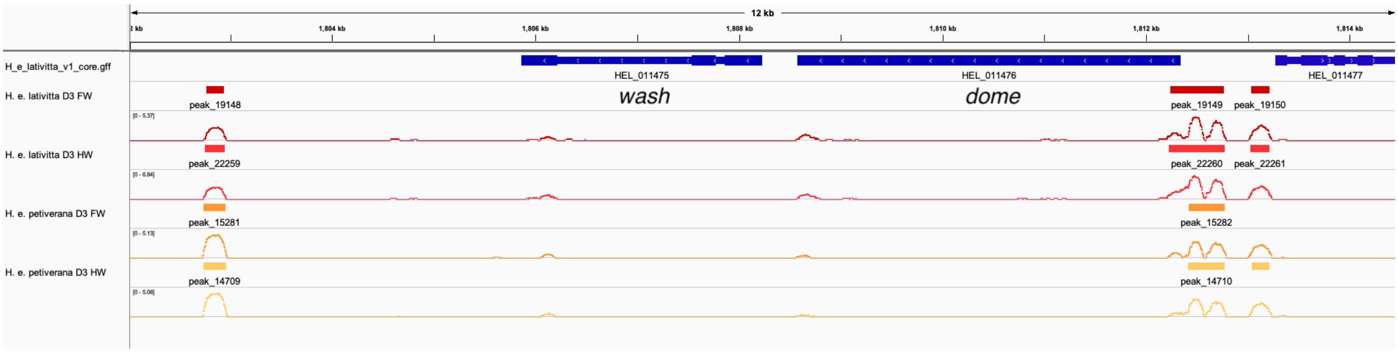


**Figure S2.2:** ATAC-sequencing analysis supports *dome/wash* bi-cistronic transcription in *Heliconius erato*. We explored whether *dome/wash* share a promoter by analysing published ATAC-sequencing data from (Lewis *et al*, 2019). We analysed samples corresponding to the forewings and hindwings of day three pupa of *H. e. lativitta* and *H. e. petiverana.* Peaks were called using Genrich (http://github.com/jsh58/Genrich/), using the parameter -j (ATAC-seq mode) with a cutoff of $q < 0.05$ ($q$ = FDR adjusted $p$-value). Peaks were visualised in IGV. Coloured blocks correspond to significant peaks and lines represent -log($q$). Peaks, indicating *cis*-regulatory activity, were observed upstream of *dome* for all samples whereas no peaks were present upstream of the start of *wash*, indicating a shared promoter for *dome/wash*.
