## Supplementary File 3 for "*Cortex cis*-regulatory switches establish scale colour identity and pattern diversity in *Heliconius*"

**Supplementary File 3:** Diagnostic SNP analysis and qPCR confirm direction of expression change in *Heliconius* populations analysed.

**Table S3.1:** RNAseq samples were genotyped relative to protein coding WGS SNPs from individuals from the source populations in Panama. Both SNPs were contained in the protein coding sequence of the gene *Cortex*. Individuals from the RNAseq experiment match the genotype of the source populations.

| **Sequence** | **Race** | **Individual** | **Informative site, scaffold 215006** | |
|---|---|---|---|---|
| | | | 1207068 | 1210502 |
| **WGS** | *melpomene* | P3_1 | TT | TT |
| | | P3_2 | TT | TT |
| | | P3_3 | TC | TC |
| **WGS** | *rosina* | ros10_1 | CC | CC |
| | | ros10_2 | CC | CC |
| | | ros10_3 | CC | CC |
| | | ros10_4 | CC | CC |
| | | ros10_5 | CC | CC |
| | | ros10_6 | CC | CC |
| | | ros10_7 | CC | CC |
| | | ros10_8 | CC | CC |
| | | ros10_9 | CC | CC |
| | | ros10_10 | CC | CC |
| **RNAseq** | *melpomene* | 49 | TT | TT |
| | | 52 | TT | TT |
| | | 54 | TT | TT |
| | | 55 | TT | TT |
| | | 56 | TT | TT |
| | | 61 | TT | TT |
| | | 62 | TT | TT |
| **RNAseq** | *rosina* | 13 | CC | CC |
| | | 14 | CC | CC |
| | | 15 | CC | CC |
| | | 16 | CC | CC |
| | | 17 | CC | CC |
| | | 24 | CC | CC |

**Table S3.2:** This was repeated for the *H. erato* samples; here, only one informative protein-coding SNP was found, in the gene *parn*. Once again, all individuals match the expected genotype.

| | | | **Informative site, scaffold 1505** |
|---|---|---|---|
| **Sequence** | Race | Individual | 2306177 |
| **WGS** | *hydara* | STRI_WOM_0039 | AA |
| | | STRI_WOM_0040 | AA |
| | | STRI_WOM_0042 | AA |
| | | STRI_WOM_0088 | AA |
| | | STRI_WOM_5193 | AA |
| | | STRI_WOM_5351 | AA |
| **WGS** | *demophoon* | Pet_ED3 | GG |
| | | Pet_ED4 | GG |
| | | Pet_ED5 | GG |
| | | Pet_ED6 | GG |
| | | STRI_WOM_0033 | GG |
| | | STRI_WOM_0082 | GG |
| | | STRI_WOM_0087 | GG |
| | | STRIWOM1284 | GG |
| | | STRIWOM5353 | GG |
| | | STRIWOM5362 | GG |
| **RNAseq** | *hydara* | 17 | AA |
| | | 25 | AA |
| | | 33 | AA |
| | | 34 | AA |
| | | 36 | AA |
| **RNAseq** | *demophoon* | A4 | GG |
| | | D2 | GG |
| | | D6 | GG |
| | | D9 | GG |
| | | C6 | GG |
| | | H3 | GG |
| | | A4 | GG |

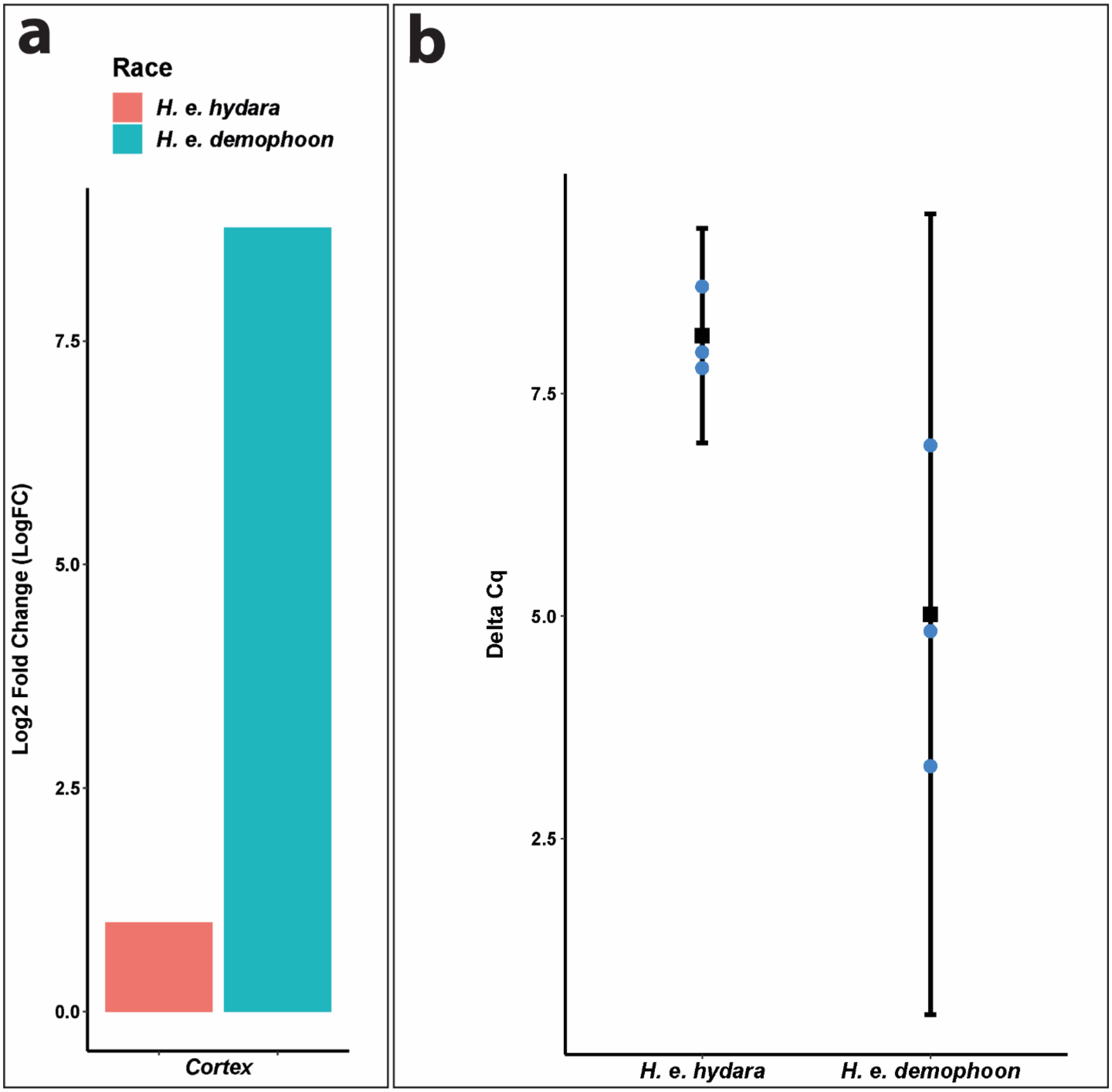


**Figure S3.1:. qPCR experiments show that cortex is expressed at a higher level in *H. e. demophoon* than in *H. e. hydara*.**

(**a**) Cortex log2 fold change relative to *H. e. hydara* using delta CT. Data were normalised against the geometric average CT of three housekeeping genes *eF1a*, *rpL3* and *polyABP*. (**b**) *Cortex* deltaCT in *H. e. hydara* and *H. e. demophoon*. Error bars represent the confidence interval for n=3 (p = 0.0444).

**Table S3.3:** Primers used for qPCR

| **Locus** | **PCR Primers (5’ > 3’)** |
|---|---|
| ***eF1α*** | AAGAATTCCCTCCCCTCGGT<br>CCACCAGCACCTTCCTTGAA |
| ***rpL3*** | AAGTCCCTTCGTGTCCACAC<br>GTGTCTGGAAGCGACCATGT |
| ***polyABP*** | CTACGCACATCTCTCGGAGC<br>CAGCTGGACGTCCACTTGTA |
| ***cortex*** | ATAGGACTGGCGAGCTGGTA<br>TTCGTACTCGGTCCAGTCCA |
