## Supplementary File 4 for "*Cortex cis*-regulatory switches establish scale colour identity and pattern diversity in *Heliconius*"

**Supplementary File 4:** DGE analysis show *cortex* and *dome/wash* are consistently differentially expressed between colour pattern races and pupal wing sections.

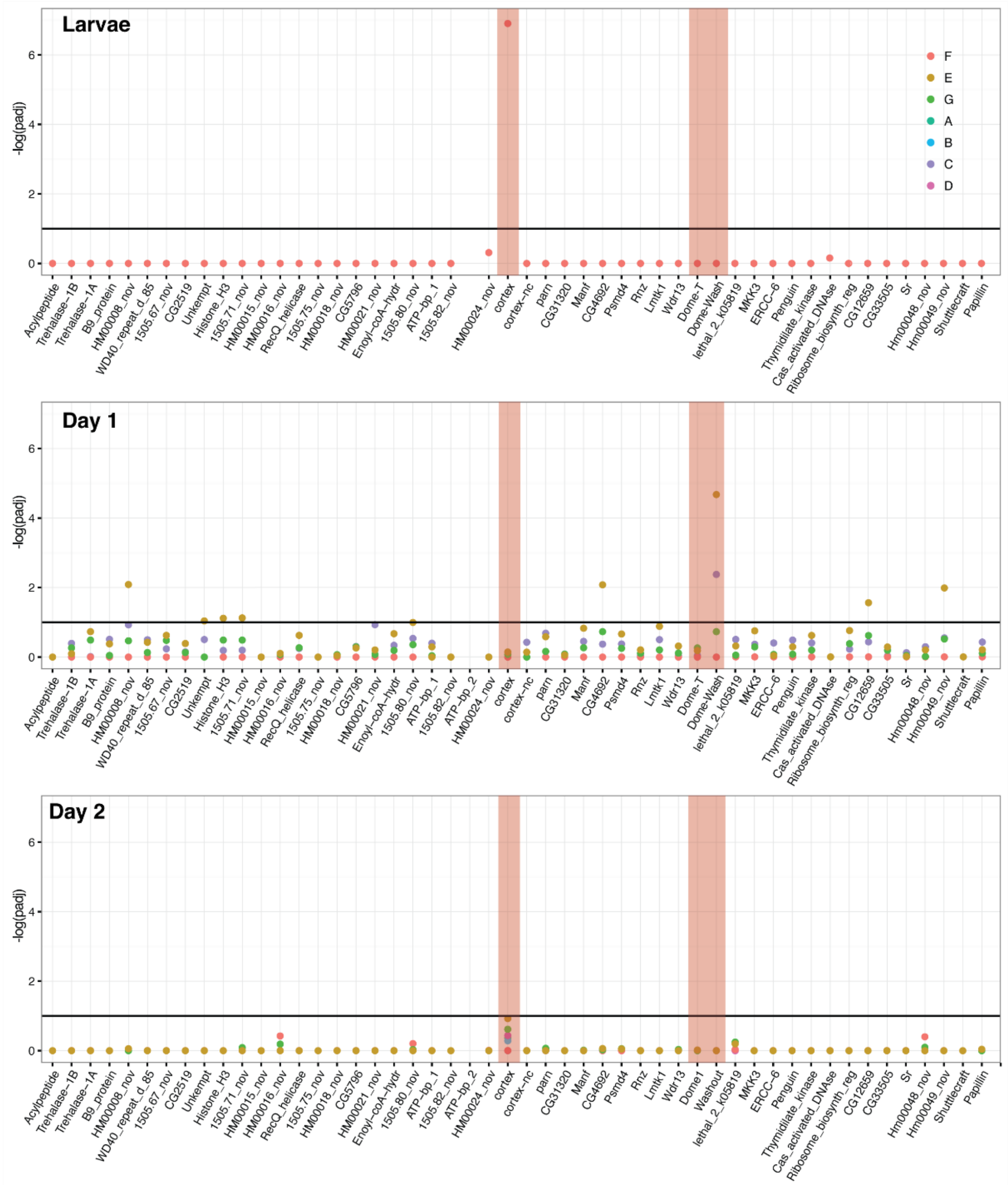


**Figure S4.1**: Differential expression across the *cortex* locus in *H. erato*, shown as the negative log of the adjusted p value (-log(*padj*) ). Top: larvae, middle: day 1 pupae, bottom, day 2 pupae. See below table for gene IDs and homology with *H. melpomene*. The red shading highlights the genes *cortex*, *dome-T* and *Dome-Wash*. The horizontal line indicates the cutoff for significance, at *padj*=0.1. Colours are used for each of the contrasts, depicted in Figure 3. In this analysis, genes were differentially expressed in contrast E and C (depictions of these contrasts are provided). E gives genes differentially regulated in black posterior compartment, and C gives genes differentially regulated in yellow anterior compartment.

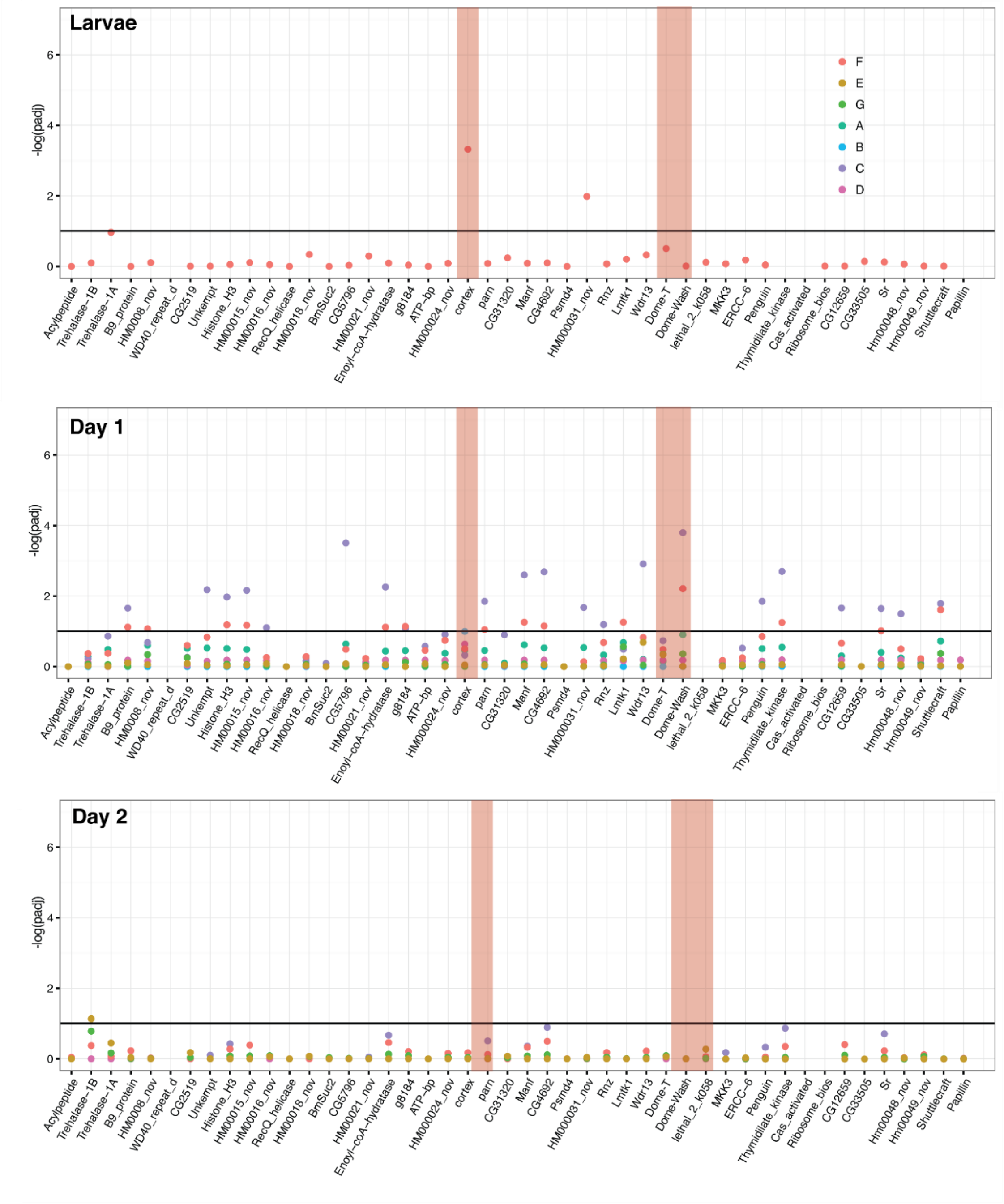


**Figure S4.2**. Differential expression across the *cortex* locus in *H. melpomene,* shown as the negative log of the adjusted p value (-log(*padj*). Top: larvae, middle: day 1 pupae, bottom, day 2 pupae. See below table for gene IDs and homology with *H. erato.* Red bars highlight the genes *cortex*, *Dome-T* and *Dome-Wash*. The horizontal line indicates the cutoff for significance, at *padj*=0.1. Colours are used for each of the contrasts, depicted in Figure S4.3. In this analysis, genes were differentially expressed in contrast E, C and F (depictions of these contrasts are provided). F is the difference between races, E gives genes differentially regulated in black posterior compartment, and C gives genes differentially regulated in yellow anterior compartment (these contrasts are depicted in cartoon form).

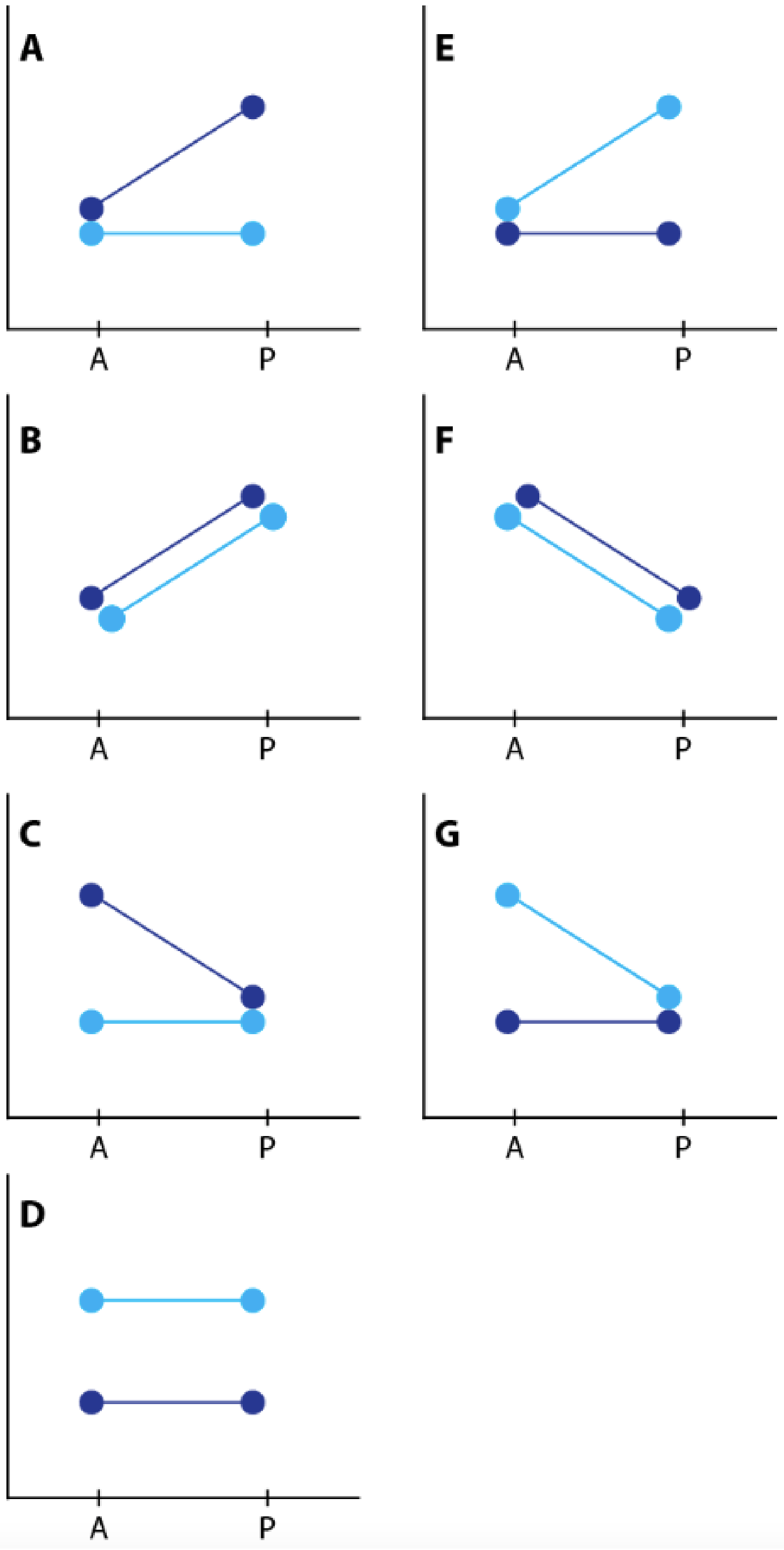


**Figure S4.3:** Depiction of contrasts. Dark blue represents the yellow races, *H. m. rosina* and *H. e. demophoon* from Panama. Light blue represents the black races, *H. m. melpomene* and *H. e. hydara* from Colombia.

**Table S4.1:** Gene IDs in the *H. melpomene* Yb locus and their corresponding IDs in the *H. erato* genome.

| Hera1 Gene ID | gene name / description | Hmel2 Gene ID |
|---|---|---|
| evm.model.Herato1505.61 | Acylpeptide | HMEL000003 |
| evm.model.Herato1505.62 | Trehalase-1B | HMEL000004 |
| evm.model.Herato1505.63 | Trehalase-1A | HMEL000006 |
| evm.model.Herato1505.64 | B9_protein | HMEL000007 |
| evm.model.Herato1505.65 | HM0008_nov | HMEL000008 |
| evm.model.Herato1505.66 | WD40_repeat_d_85 | HMEL000010 |
| evm.model.Herato1505.67 | | |
| evm.model.Herato1505.68 | CG2519 | HMEL000012 |
| evm.model.Herato1505.69 | Unkempt | HMEL000013 |
| evm.model.Herato1505.70 | Histone_H3 | HMEL000014 |
| evm.model.Herato1505.71 | | |
| evm.model.Herato1505.72 | HM00015_nov | HMEL000015 |
| evm.model.Herato1505.73 | HM00016_nov | HMEL000016 |
| evm.model.Herato1505.74 | RecQ_helicase | HMEL000017 |
| evm.model.Herato1505.75 | | |
| evm.model.Herato1505.76 | HM00018_nov | HMEL000018 |
| *****(1505.35!)***** | BmSuc2 | HMEL000019 |
| evm.model.Herato1505.77 | CG5796 | HMEL000020 |
| evm.model.Herato1505.78 | HM00021_nov | HMEL000021 |
| evm.model.Herato1505.79 | Enoyl-coA-hydratase | HMEL000022 |
| evm.model.Herato1505.80 | | |
| evm.model.Herato1505.81 | ATP-bp | HMEL002023 |
| evm.model.Herato1505.82 | | |
| evm.model.Herato1505.83 | HMEL002023 | |
| evm.model.Herato1505.84 | HM00024_nov | HM00024_nov |
| | | g8186 |
| evm.model.Herato1505.85 | Cortex | HMEL000025 |
| evm.model.Herato1505.86 | parn | HMEL000026 |
| evm.model.Herato1505.87 | CG31320 | HMEL000027 |
| evm.model.Herato1505.88 | Manf | HMEL000028 |
| evm.model.Herato1505.89 | CG4692 | HMEL000029 |
| evm.model.Herato1505.90 | Psmd4 | HMEL000030 |
| | HM000031_nov | HMEL000031 |
| evm.model.Herato1505.91 | Rnz | HMEL000032 |
| evm.model.Herato1505.92 | Lmtk1 | HMEL000033 |
| evm.model.Herato1505.93 | Wdr13 | g8202 |
| evm.model.Herato1505.94 | Dome1 | HMEL013472 |
| evm.model.Herato1505.95 | Washout | HMEL000036 |
| evm.model.Herato1505.96 | Dome2 | HMEL000037 |
| evm.model.Herato1505.97 | lethal_2_k05819 | g8206 |
| evm.model.Herato1505.98 | MKK3 | HMEL000039 |

| | | |
|---|---|---|
| evm.model.Herato1505.99 | ERCC-6 | HMEL000040 |
| evm.model.Herato1505.100 | Penguin | HMEL000041 |
| evm.model.Herato1505.101 | Thymidilate_kinase | HMEL000042 |
| evm.model.Herato1505.102 | Cas_activated_DNAse | HMEL000043 |
| evm.model.Herato1505.103 | Ribosome_biosynth_r | HMEL000044 |
| evm.model.Herato1505.104 | CG12659 | HMEL000045 |
| evm.model.Herato1505.105 | CG33505 | HMEL000046 |
| evm.model.Herato1505.106 | Sr | HMEL000047 |
| evm.model.Herato1505.107 | Hm00048_nov | HMEL000048 |
| evm.model.Herato1505.108 | Hm00049_nov | HMEL000049 |
| evm.model.Herato1505.109 | | HMEL000050 |
| | Papillin | HMEL000051 |
| evm.model.Herato1505.110 | HM000052_nov | HMEL000052 |
