## Supplementary File 5 for "*Cortex cis*-regulatory switches establish scale colour identity and pattern diversity in *Heliconius*"

**Supplementary File 5**: Cortex is a derived and insect-specific derivative of Cdc20.

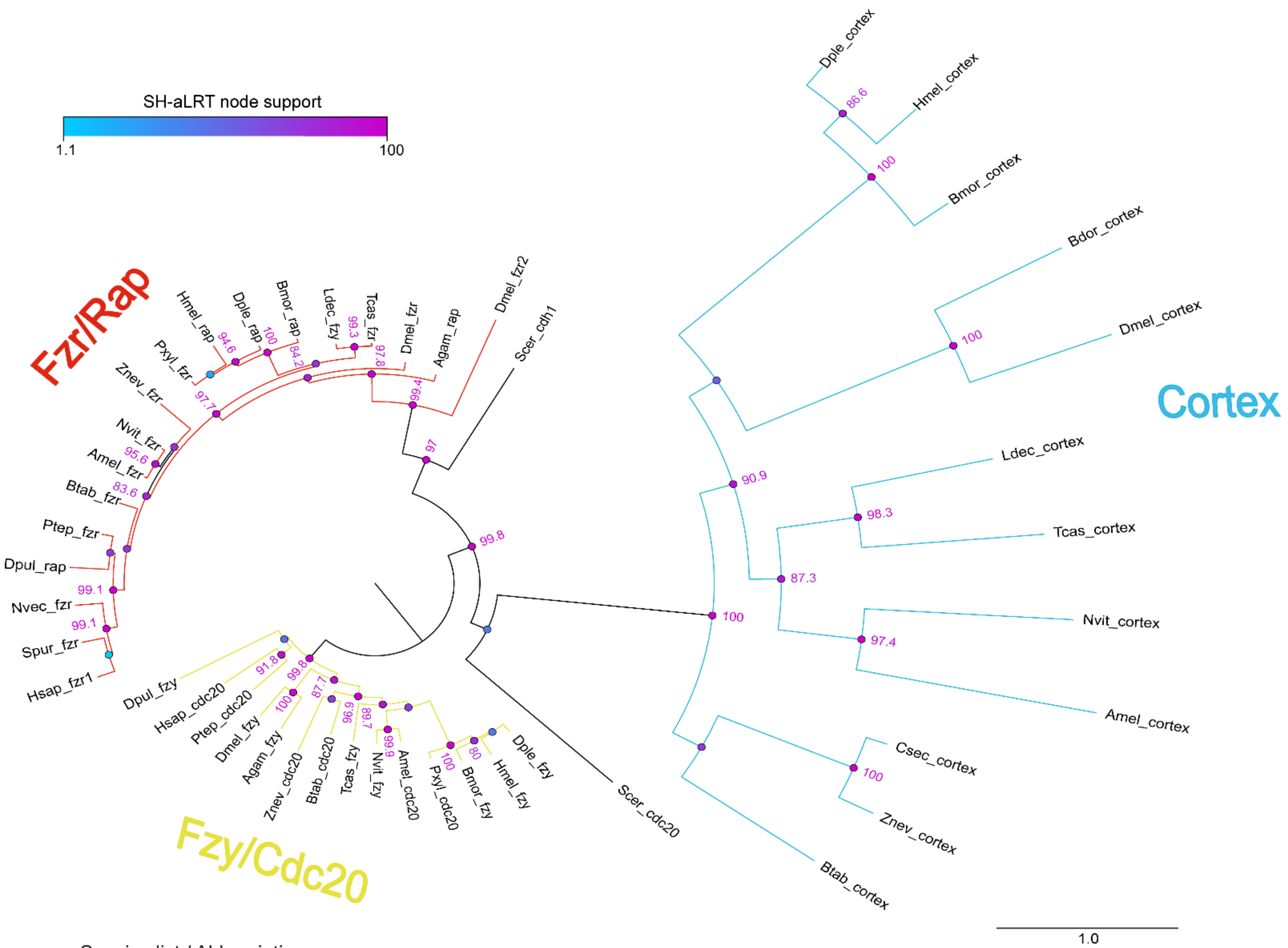


Species list / Abbreviations

| | |
|---|---|
| Saccharomyces cerevisiae | Fungi |
| Nematostella vectensis | Cnidaria |
| Aplysia californica | Mollusca |
| Parasteatoda tepidariorum | Arachnida |
| Strongylocentrotus purpuratus | Echinodermata |
| Homo sapiens | Vertebrata |

| | | | |
|---|---|---|---|
| Anopheles gambiae | Diptera | Bombyx mori | Lepidoptera |
| Drosophila melanogaster | Diptera | Danaus plexippus | Lepidoptera |
| Bactrocera dorsalis | Diptera | Heliconius melpomene | Lepidoptera |
| Apis mellifera | Hymenoptera | Plutella xylostella | Lepidoptera |
| Nasonia vitripennis | Hymenoptera | | |
| Leptinotarsa decemlineata | Coleoptera | | |
| Tribolium castaneum | Coleoptera | | |
| Bemisia tabaci | Hemiptera | | |
| Zootermopsis nevadensis | Blattodea | | |
| Cryptotermes secundus | Blattodea | | Neoptera |

**Supplementary File 5 - Figure S5.1**. Phylogenetic analysis of the cdc20/cdh1 family reveals Cortex is a derived and insect-specific derivative of cdc20. Full-length protein homologs retrieved from TBLASTN searches were used to generate a curated alignment with MAFFT/Guidance2 with a column threshold of 0.5. TBLASTN searches against arthropod genome and transcriptome NCBI repositories did not recover Cortex homologues outside of the Neoptera lineage. The maximum-likelihood tree was constructed with W-IQ-TREE with the “Auto” function to find a best-fit model of substitution. Colour circles indicate the scores of SH-like approximate likelihood ratio tests (SH-aLRT) computed over 1,000 replicates, with numeric values for scores > 80. Scale bar indicates amino-acid substitutions per site. Abbreviations: fzy, fizzy ; fzr, fizzy-related ; rap, retina aberrant pattern (syn. fzr).

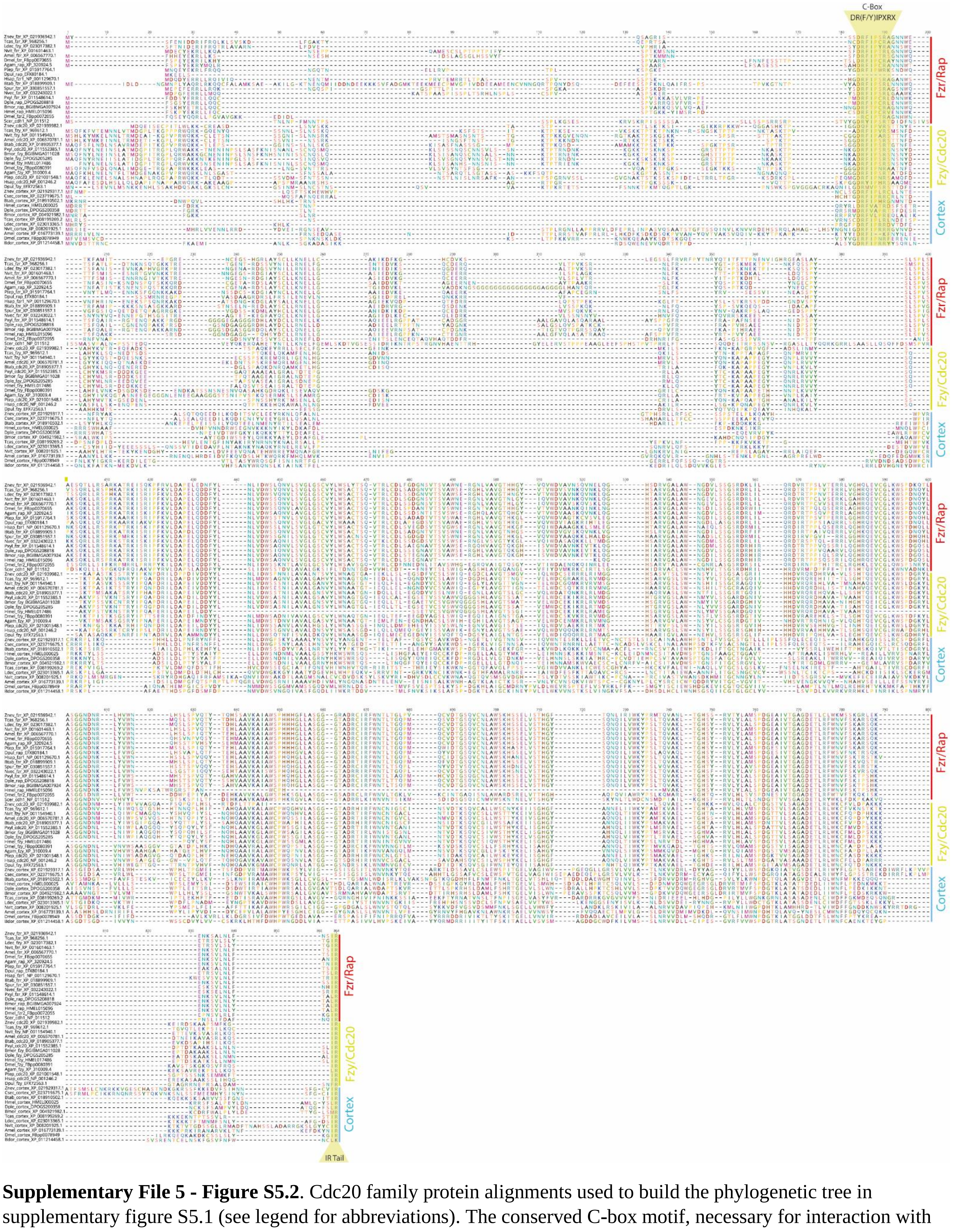


**Supplementary File 5 - Figure S5.2**. Cdc20 family protein alignments used to build the phylogenetic tree in supplementary figure S5.1 (see legend for abbreviations). The conserved C-box motif, necessary for interaction with the APC/C and the IR are highlighted.
