## Supplementary File 6 for "*Cortex cis*-regulatory switches establish scale colour identity and pattern diversity in *Heliconius*"

**Supplementary File 6:** Distal expression of *cortex* in *Heliconius* 5th instar imaginal discs.

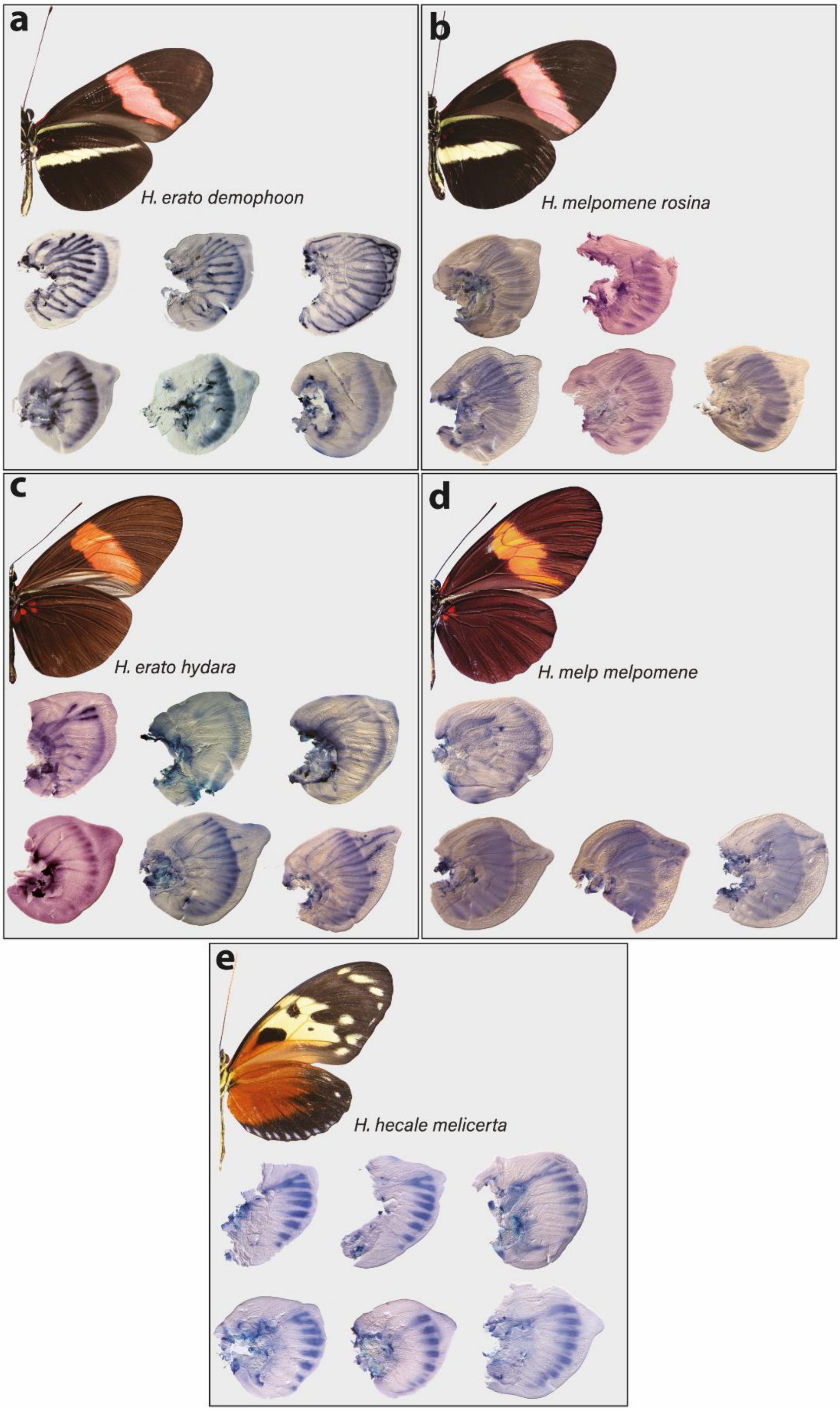


**Figure S6:** In *H. erato*, *cortex* expression is strongest at the distal end of the wing throughout 5th instar development, with stronger intervein expression in *H. erato hydara*. In *H. melpomene*, *cortex* expression extends further proximally, with expression seen throughout the wing in *H. melpoemene melpomene*.
