## Supplementary File 7 for "*Cortex cis*-regulatory switches establish scale colour identity and pattern diversity in *Heliconius*"

**Supplementary File 7:** CRISPR mutagenesis confirmed through sanger sequencing

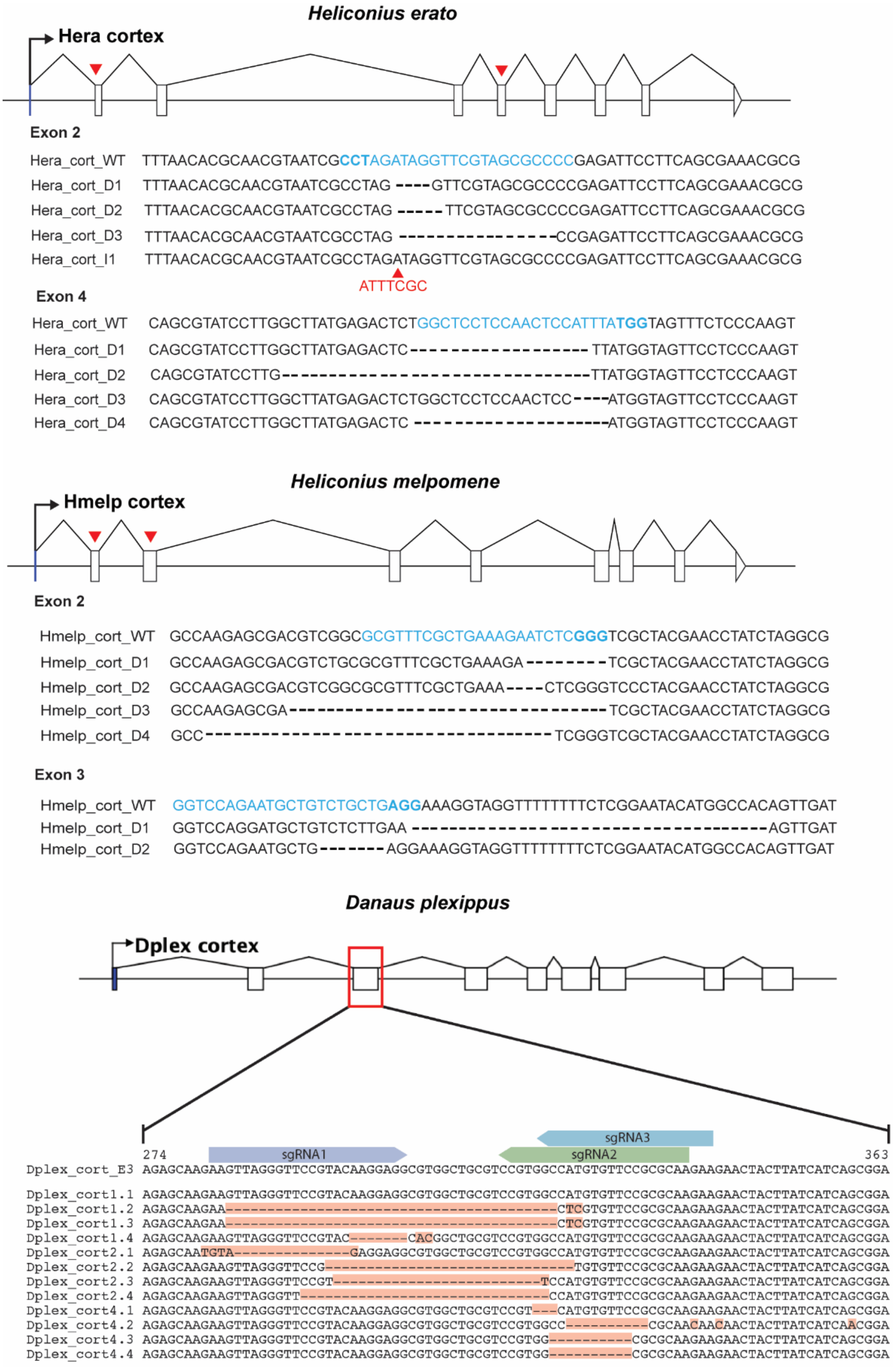


**Figure S7.1:** Gene models for cortex are shown for both *H. erato*, *H. melpomene* and *D. plexippus*. Red arrows indicate positions of sgRNAs that were genotyped in CRISPR experiments. Recovered sequences showing evidence of editing as a result of CRISPR mutagenesis are shown. Target sequences are shown in blue and PAM site highlighted in bold. For *H. erato* exon 2, we recovered a sequence containing a 7bp insertion (indicated with red arrowhead, I2).

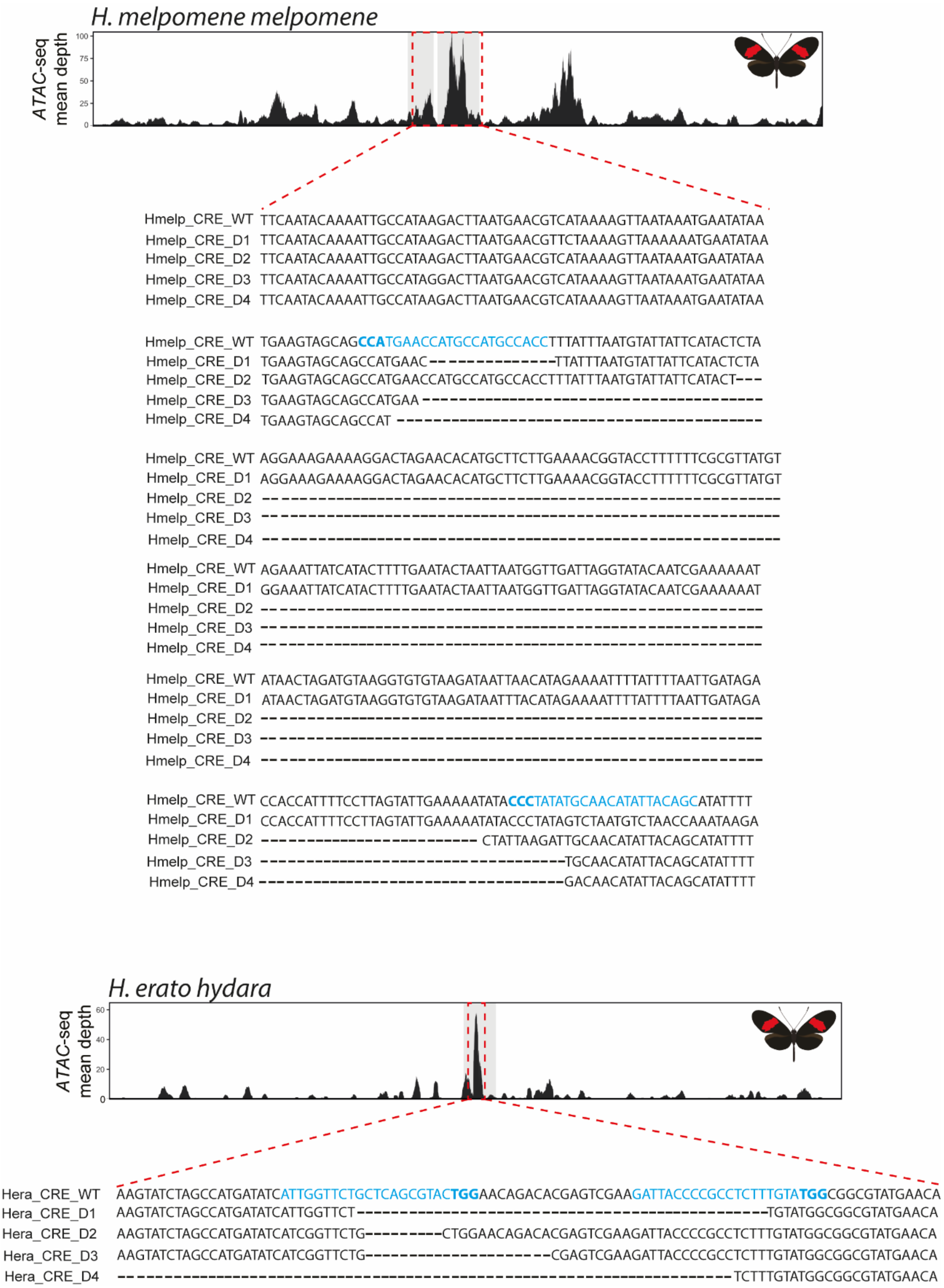


**Figure S7.2:** Genotyping at the *H. melpomene* and *H. erato* CRE confirms CRISPR induced deletions. Targeted ATAC-seq peak shown above (red dotted rectangle), as well as alignment between wild-type and recovered deletions.  Guides highlighted in blue with PAM sequence in bold.
