## Supplementary File 9 for "*Cortex cis*-regulatory switches establish scale colour identity and pattern diversity in *Heliconius*"

**Supplementary File 9:** Mutant panels showing replicates of *cortex* mutants for each species/race.

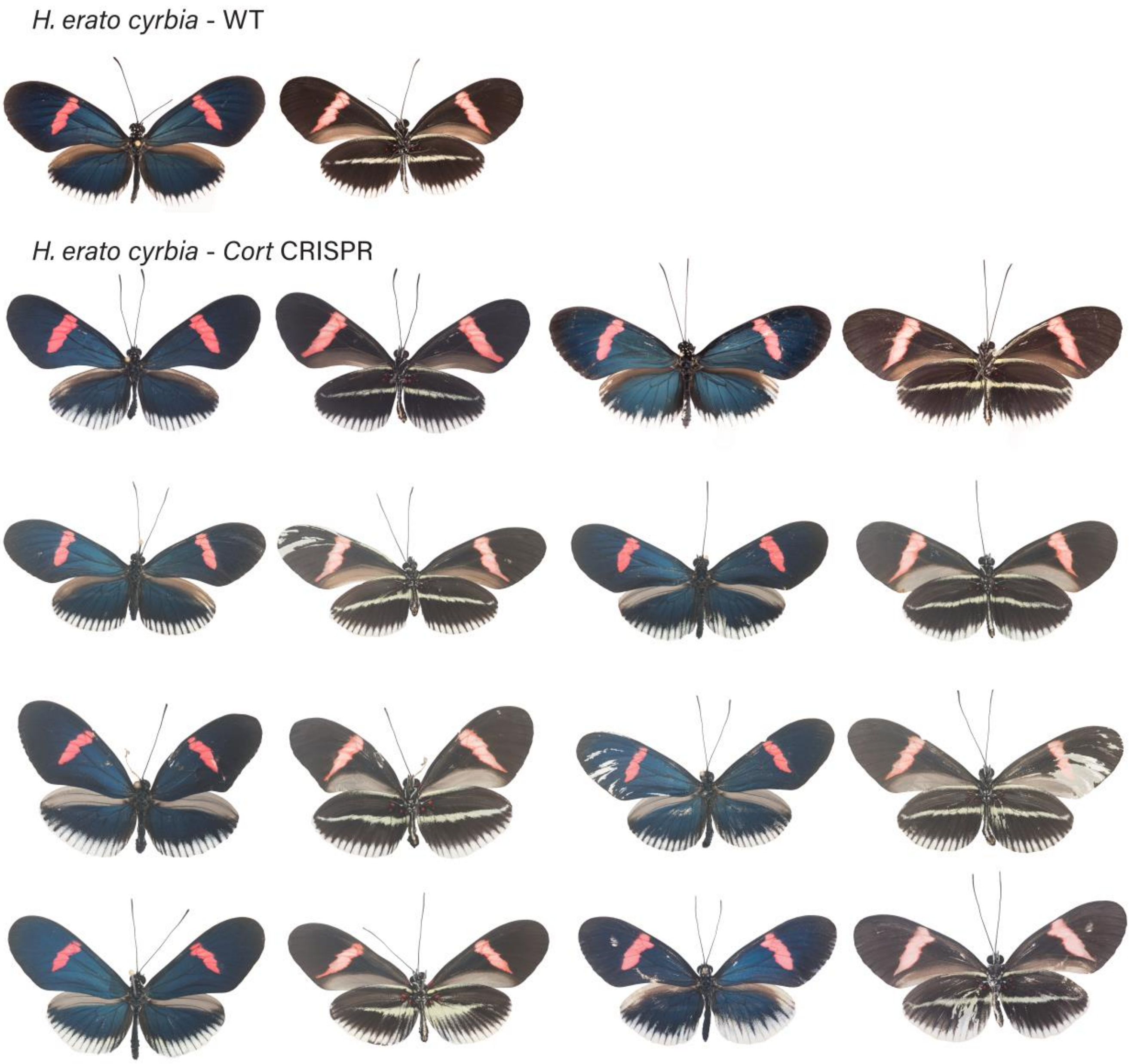


**Figure S9.1:** *H. erato cyrbia* wild-type (WT), alongside *cortex* mKO individuals recovered in CRISPR experiments. Dorsal and ventral sides shown for each mutant.

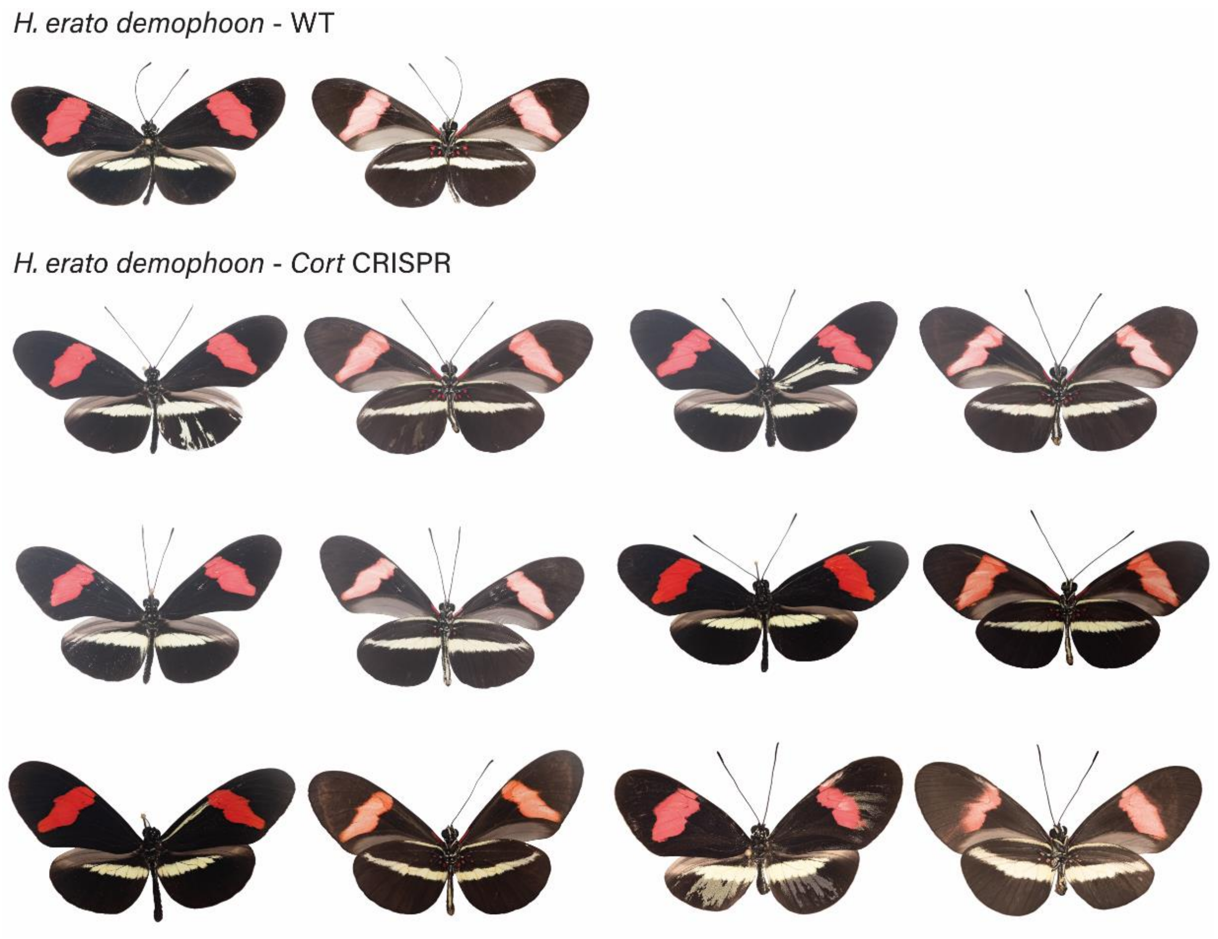


**Figure S9.2:** *H. erato demophoon* wild-type (WT), alongside *cortex* mKO individuals recovered in CRISPR experiments. Dorsal and ventral sides shown for each mutant.

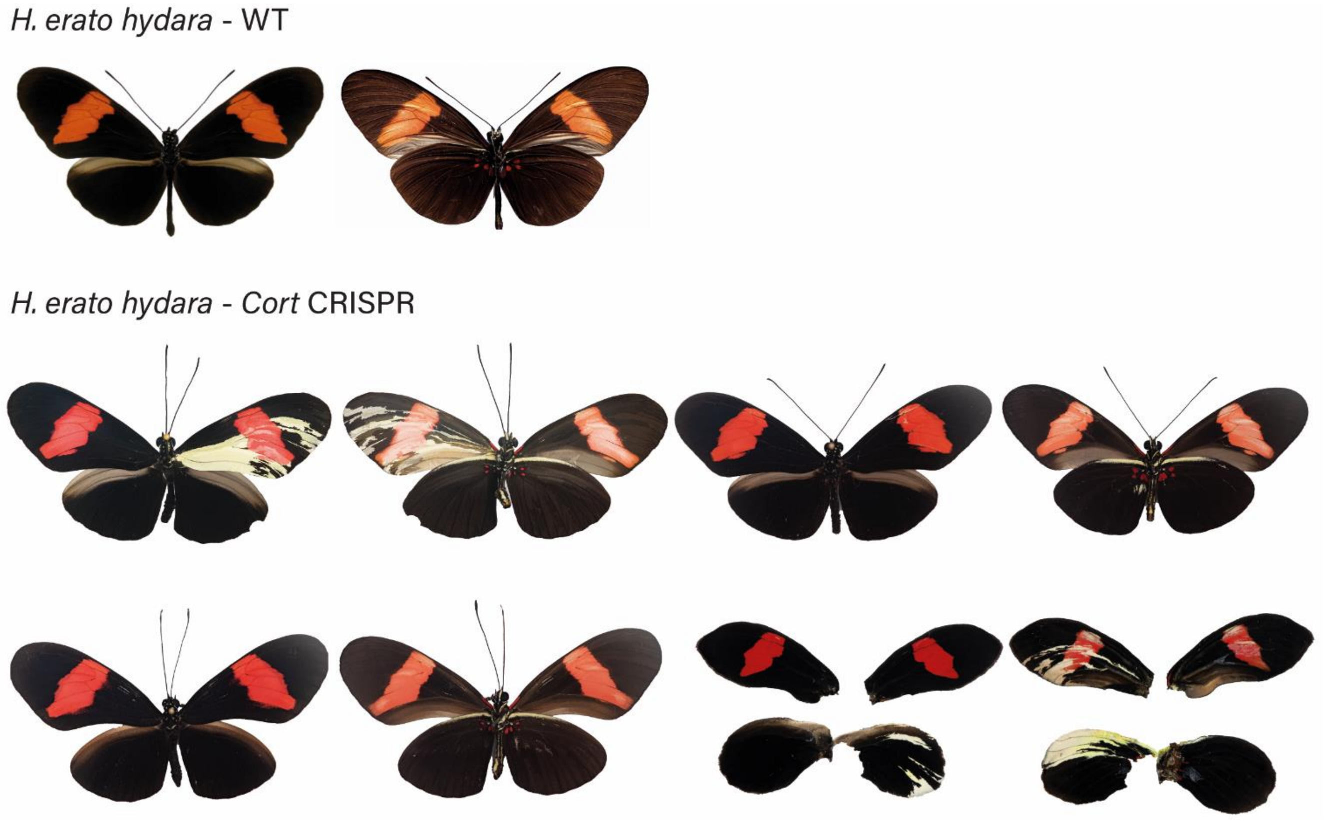


**Figure S9.3:** *H. erato hydara* wild-type (WT), alongside *cortex* mKO individuals recovered in CRISPR experiments. Dorsal and ventral sides shown for each mutant.

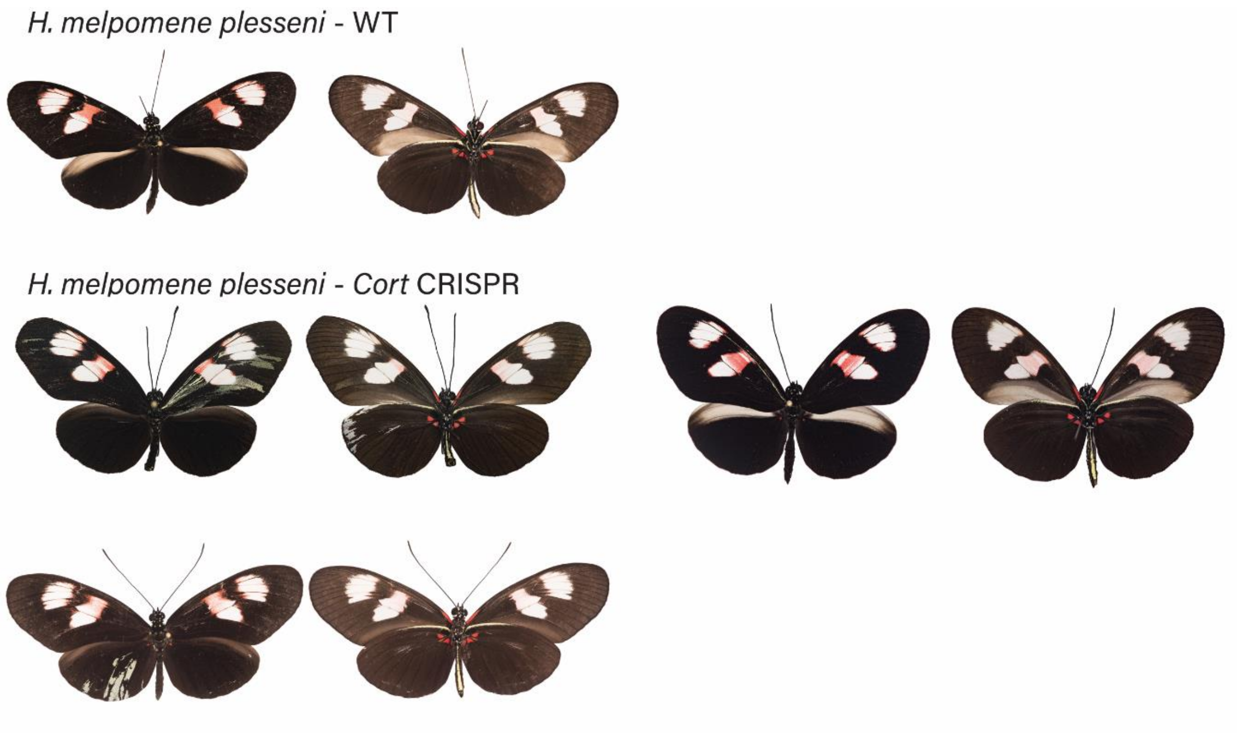


**Figure S9.4:** *H. melpomene plesseni* wild-type (WT), alongside *cortex* mKO individuals recovered in CRISPR experiments. Dorsal and ventral sides shown for each mutant.

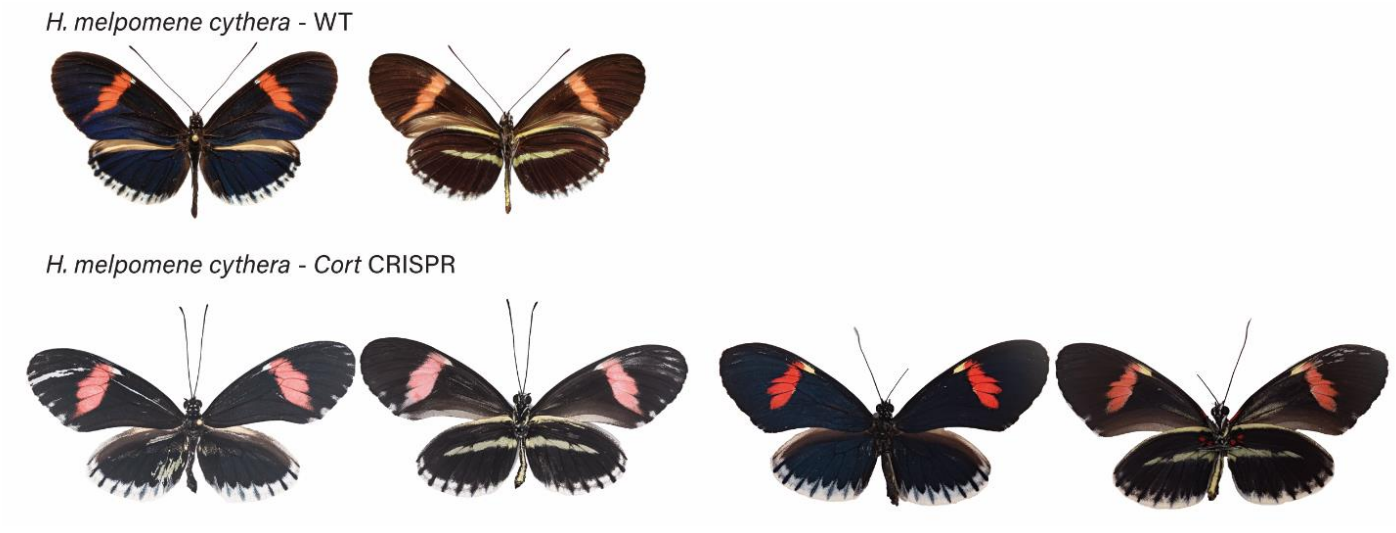


**Figure S9.5:** *H. melpomene cythera* wild-type (WT), alongside *cortex* mKO individuals recovered in CRISPR experiments. Dorsal and ventral sides shown for each mutant.

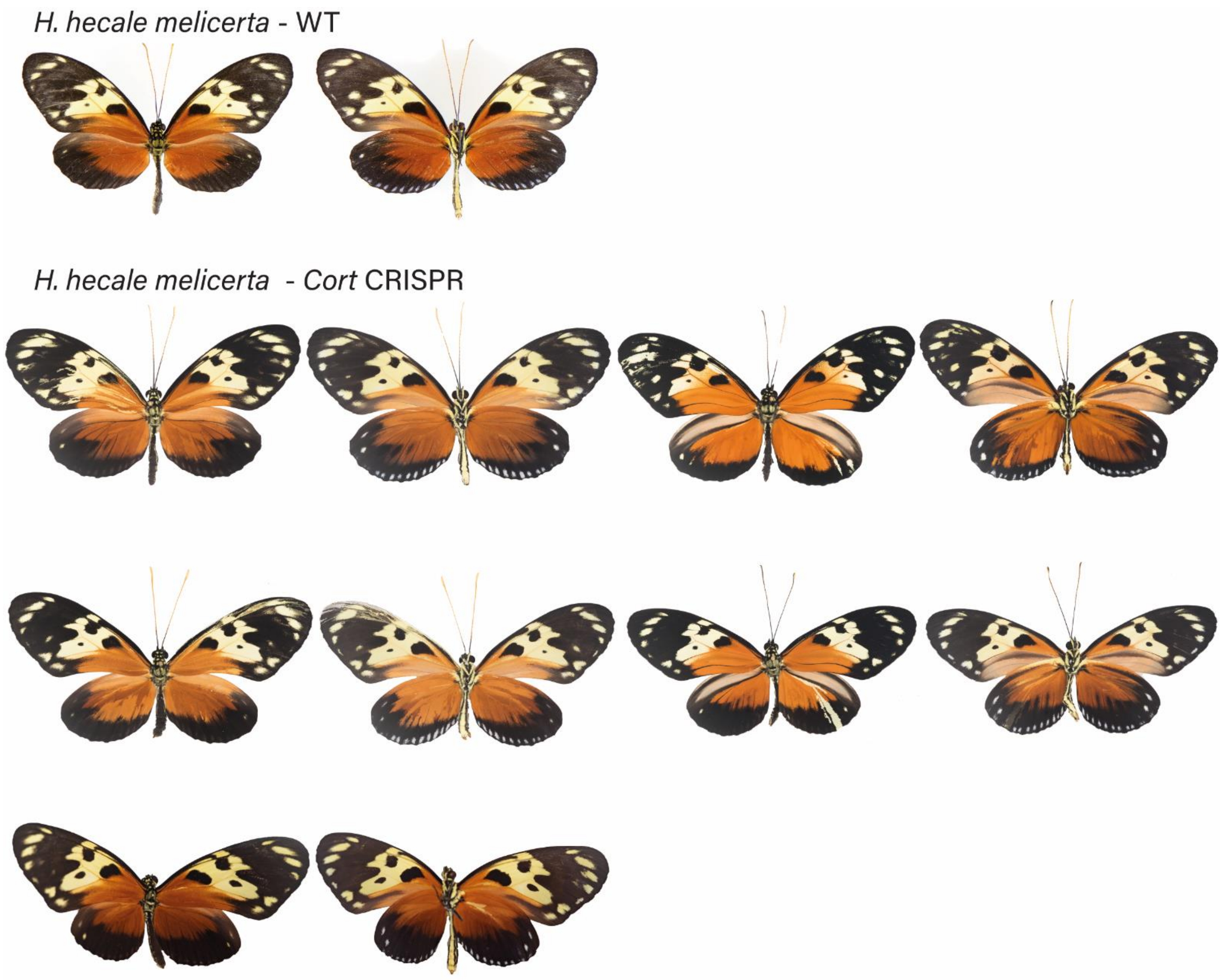


**Figure S9.6:** *H. hecale melicerta* wild-type (WT), alongside *cortex* mKO individuals recovered in CRISPR experiments. Dorsal and ventral sides shown for each mutant.

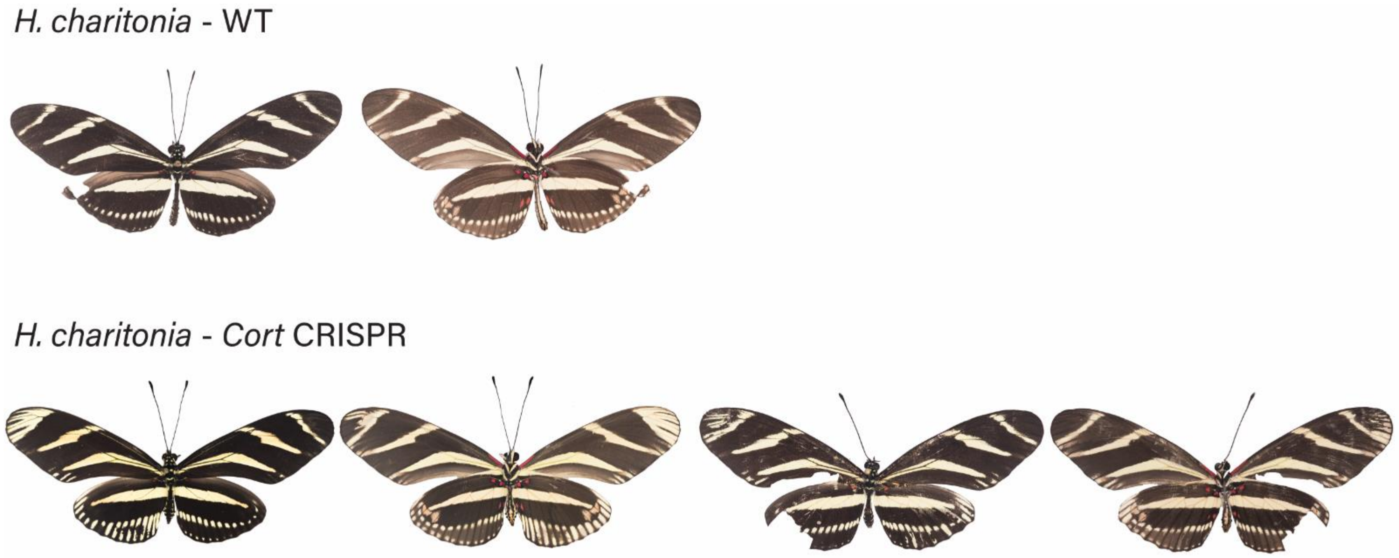


**Figure S9.7:** *H. charithonia* wild-type (WT), alongside *cortex* mKO individuals recovered in CRISPR experiments. Dorsal and ventral sides shown for each mutant.

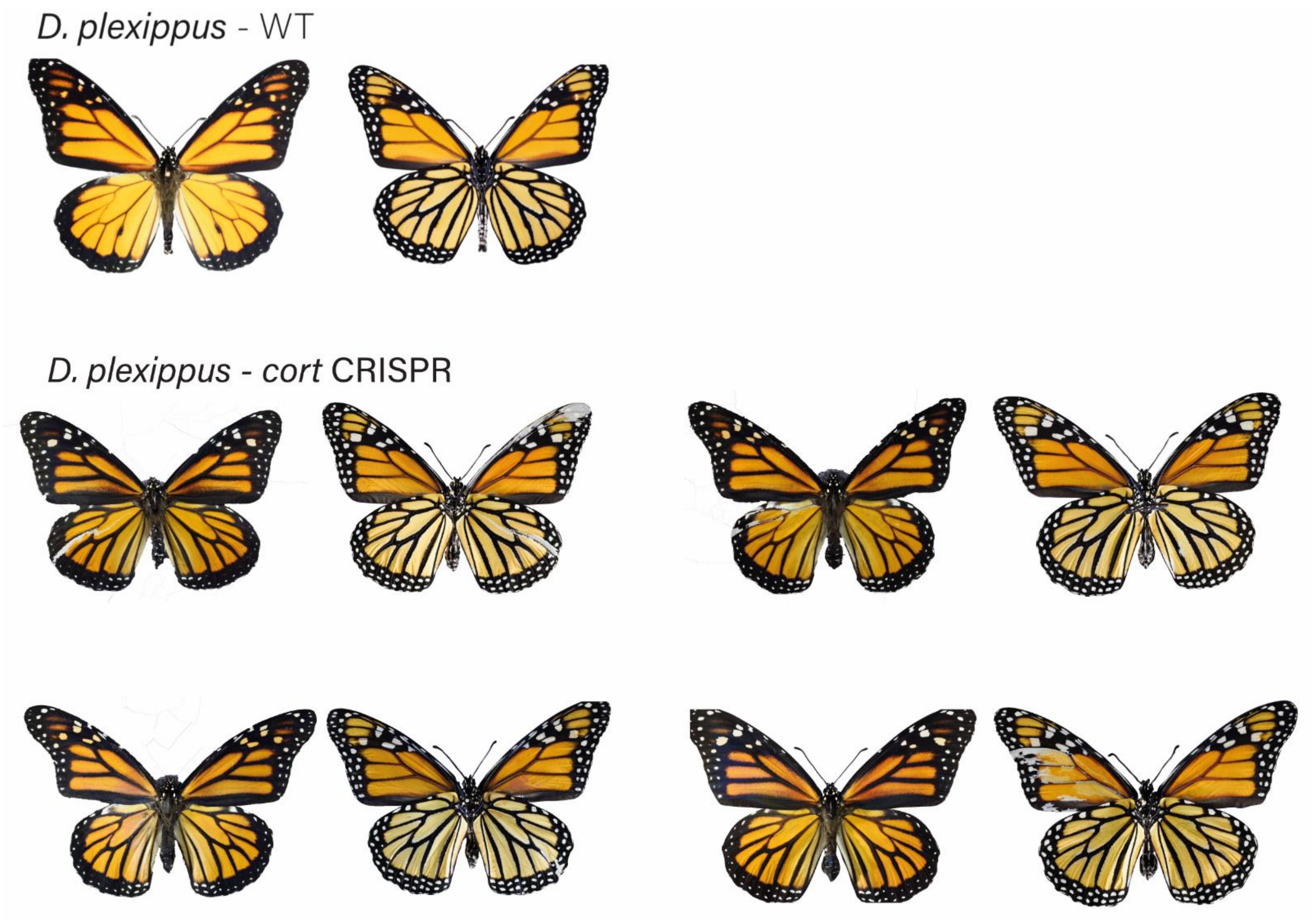


**Figure S9.8:** *Danaus plexippus* wild-type (WT), alongside *cortex* CRE mKO individuals recovered in CRISPR experiments. Dorsal and ventral sides shown for each mutant.

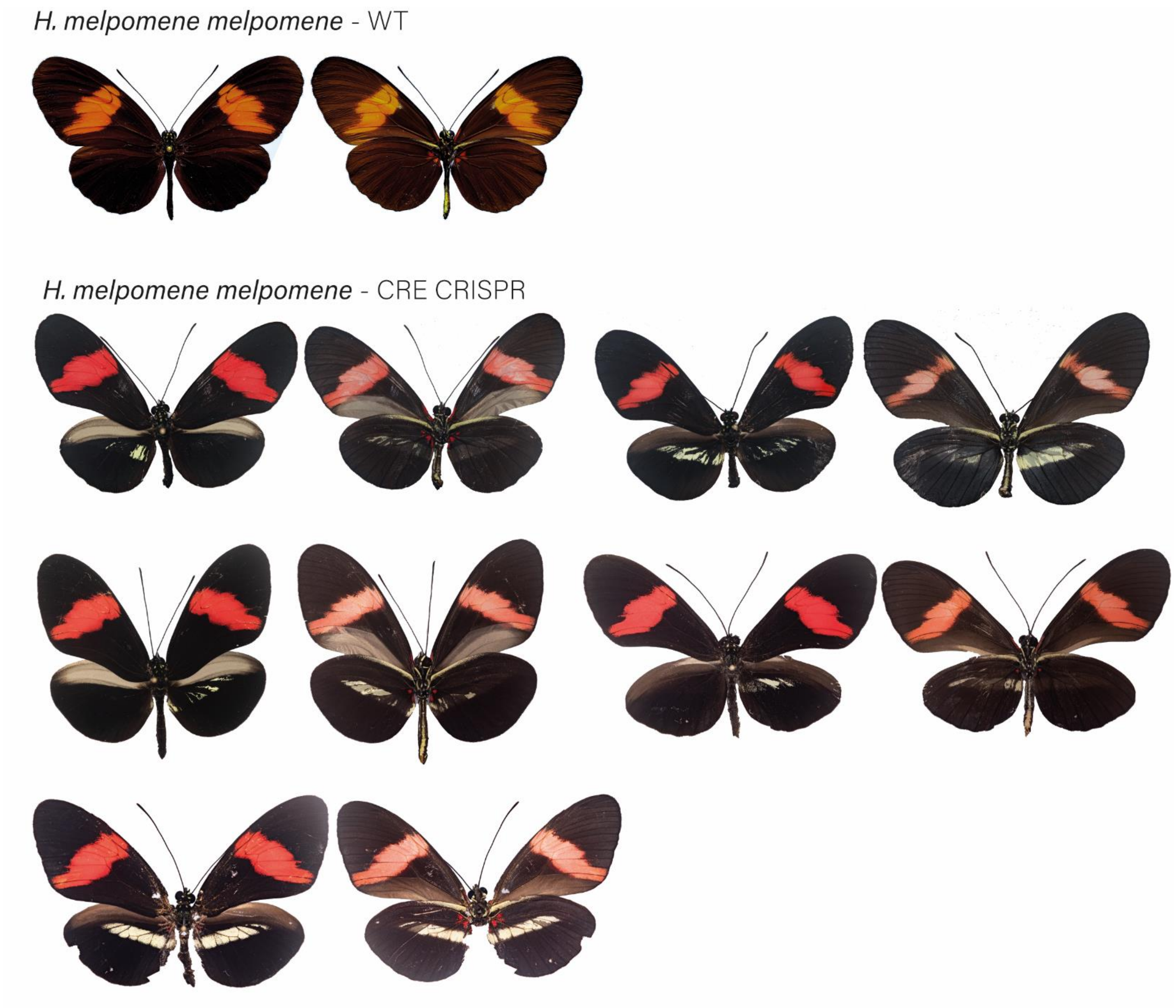


**Figure S9.9:** *H. melpomene melpomene* wild-type (WT), alongside *cortex* CRE mKO individuals recovered in CRISPR experiments. Dorsal and ventral sides shown for each mutant.

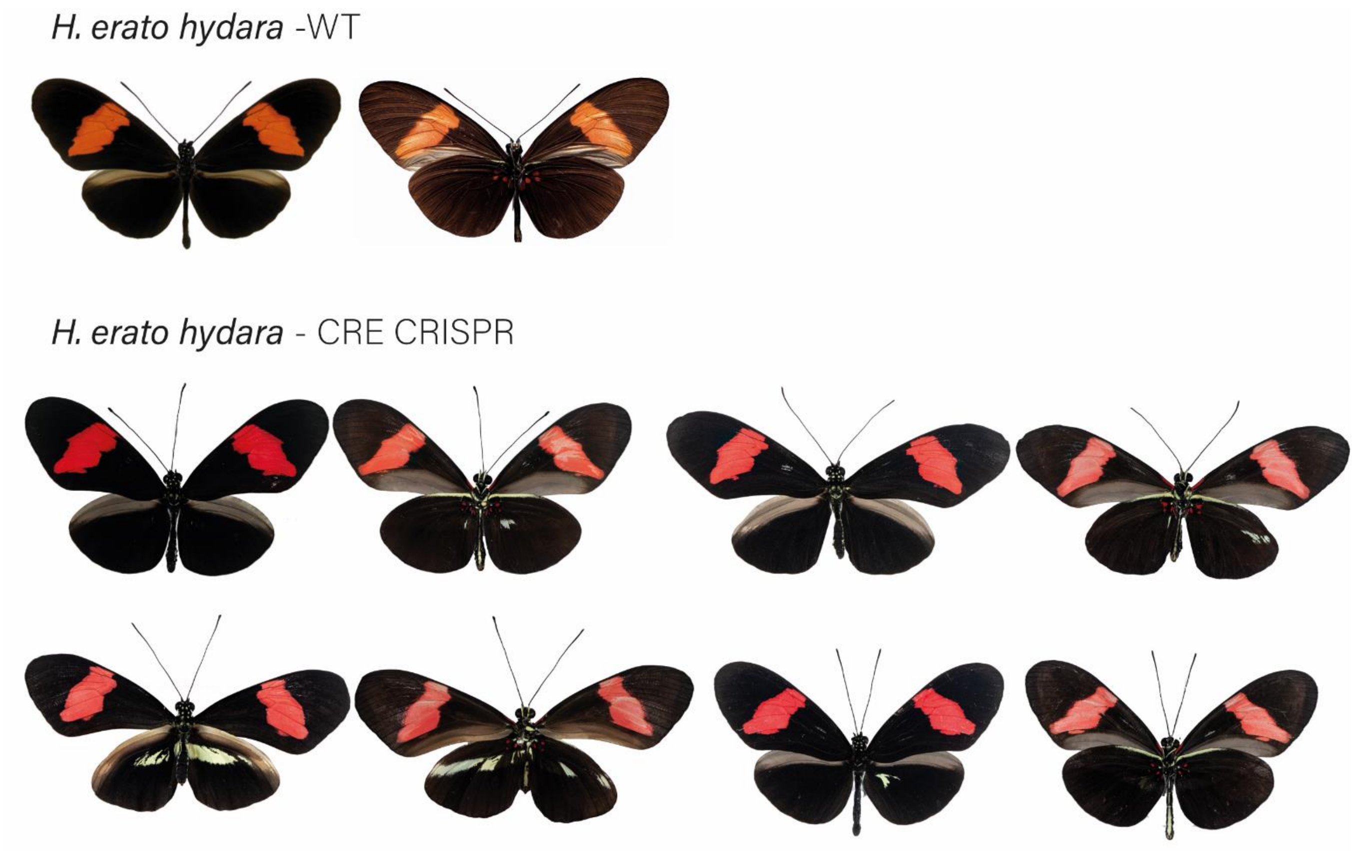


**Figure S9.10:** *H. erato hydara* wild-type (WT), alongside *cortex* CRE mKO individuals recovered in CRISPR experiments. Dorsal and ventral sides shown for each mutant.
