## Supplementary File 10 for "*Cortex cis*-regulatory switches establish scale colour identity and pattern diversity in *Heliconius*"

**Supplementary File 10**: Broad effects of *cortex* across *Heliconius* wings.

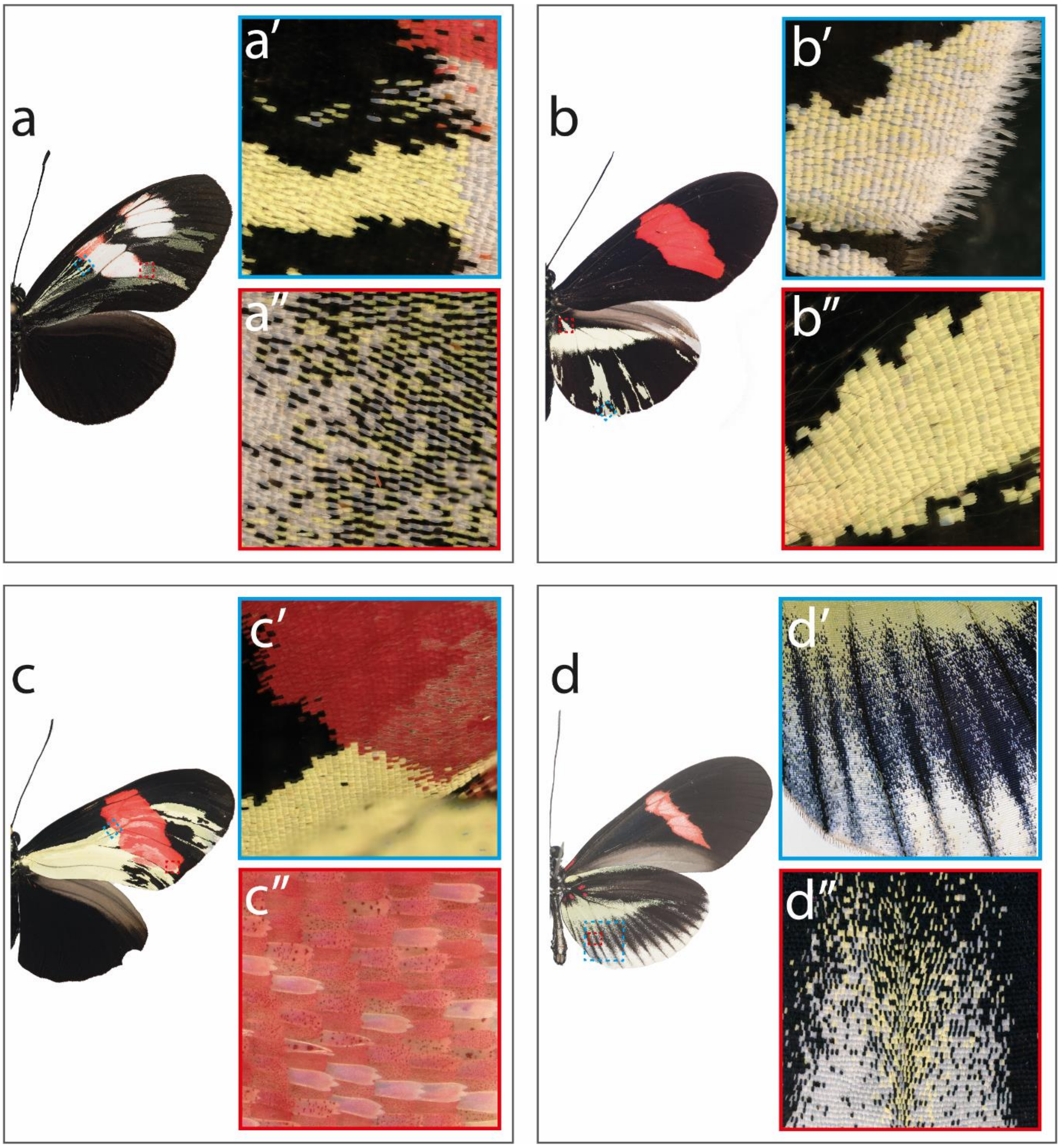


**Figure S10:** Mutant close-ups illustrating variety of effects caused by *cortex* mKO. (**a**) *H. melpomene plesseni* showing mutant clones across proximal forewing (**a'**) as we as more distal areas, where cover scales are more affected (**a''**). Several individuals show this effect (see supplementary file 7). (**b**) *H. erato demophoon* showing cortex mKO effects on elongated border scales (**b'**), and scale located anterior to yellow bar element (**b''**). (**c**) *H. erato hydara* showing clones extending into forewing red band (**c'**) and asymmetric deposition of red pigment across the affected red band region (**c''**). (**d**) *H. erato cyrbia* illustrates positional effect of cort mKO where posterior hindwing scales shift to white, while anterior scales shift to yellow (**d'**and **d''**).
