## Supplementary File 11 for "*Cortex cis*-regulatory switches establish scale colour identity and pattern diversity in *Heliconius*"

**Supplementary File 11**: **Cortex protein localises across the pupal wings of *H. erato demophoon*.**

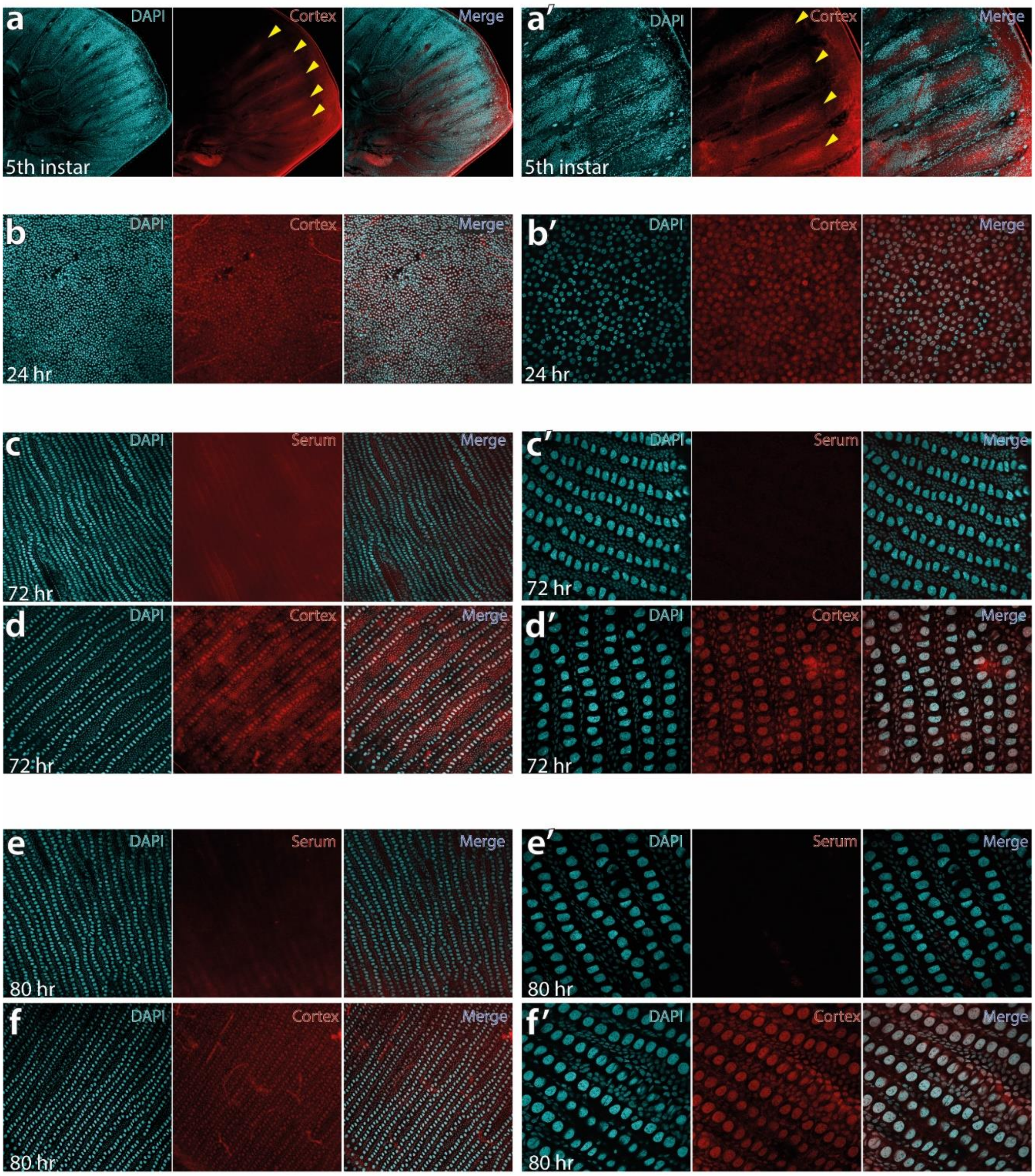


**Figure S11: Cortex antibody stainings for each sampled stage.**

Fifth instar larval wing disc showing intervein localisation of Cortex protein at 10x (**a**) and 30x magnification of the same area (a'). Pupal wings sampled at 24hr post pupation at 20x maginification (**b**) and 60x magnification (**b'**) show nuclear localisation of Cortex protein. A this stage, the hexagonal organisation of future scale cells is apparent, with the middle cell eventually differentiating into the scale cell. Pupal wings were stained with a negative control using the pre-immune serum at 72hr, shown at 20x (**c**) and 60x (**c'**), reveal no evidence of nuclear signal. Using the same confocal settings with Cortex antibody at 72hr reveals nuclear localisation at 30x magnification (**d**) and 60x magnification (**d'**). The nuclear stainig is also absent in control serum samples at 80hr post pupation (**e** and **e'**), with nuclear localisation persisiting in cortex antibody incubated samples at 80hr post pupation (**f** and **f'**).
