## Supplementary File 12 for "*Cortex cis*-regulatory switches establish scale colour identity and pattern diversity in *Heliconius*"

**Supplementary File 12:** The *cortex* locus is characterised by many accessible chromatin peaks as revealed by *ATAC-seq*

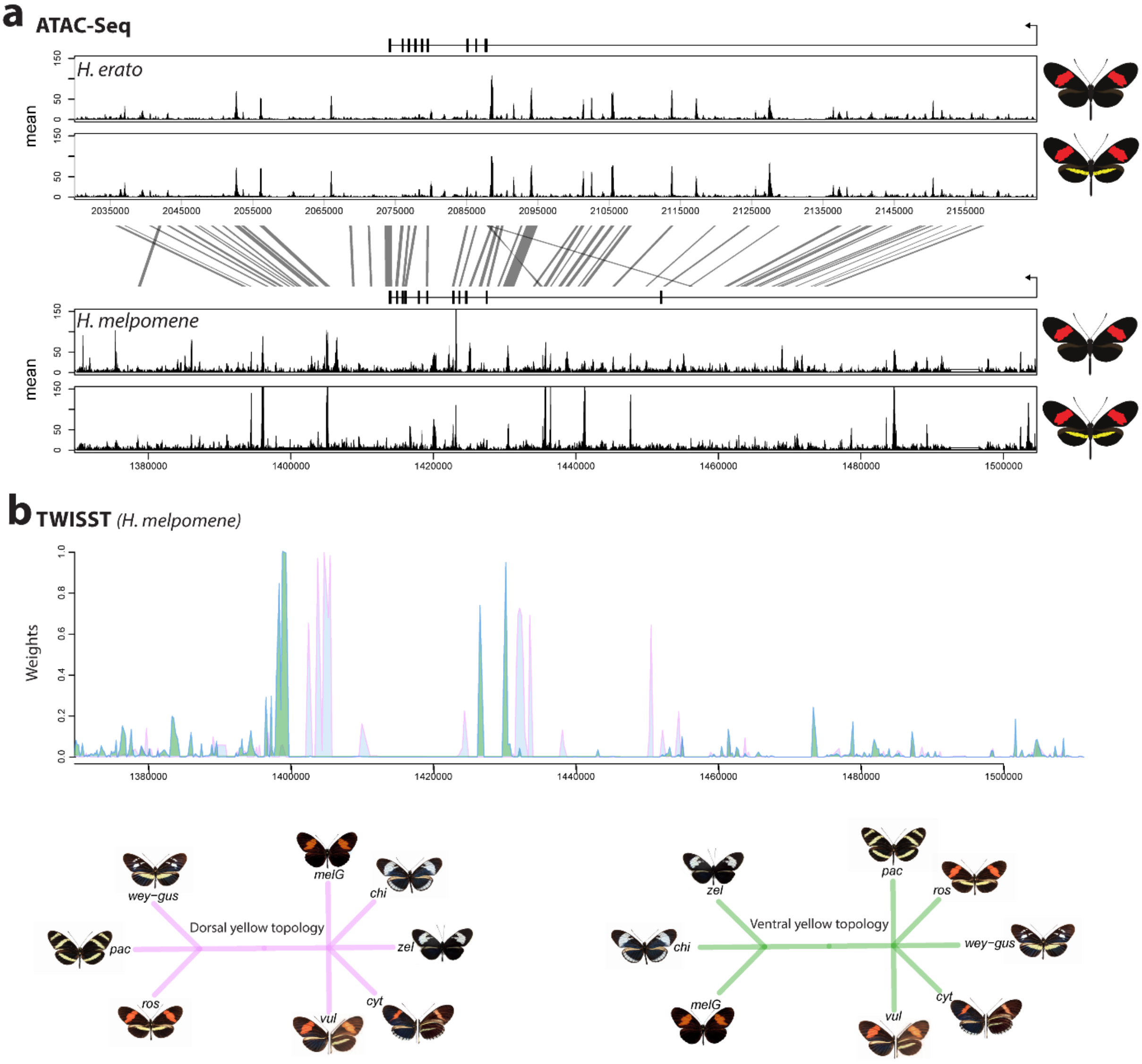


**Figure S.12:** Mean sequence depth for *ATAC-Seq* traces in *H.erato* (top) and *H. melpomene* (bottom) in 5$^{th}$ instar caterpillar hindwings in yellow banded and black morphs. The gene model for *cortex* is shown above the traces (black rectangles are coding exons, lines are introns, direction of transcription is indicated by an arrow). 95% sequence identity between *H. melpomene* and *H.erato* is indicated by grey lines. **(b)** Twisst analysis results showing high support for the presence of a yellow bar in *H.melpomene* populations with a ventral (ventral topology) or dorsal (dorsal topology) yellow bar. Tree topologies used to calculate weightings are shown. Abbreviations for twisst morphs: wey-gus = *H. cydno weymeri-gustavi*, chi = *H. cydno chioneus*, zel = *H. cydno zelinde*, pac = *H. pachinus*, ros = *H. melpomene rosina*, vul = *H. melpomene vulcanus*, cyt = *H. melpomene cythera*, melG = *H. melpomene melpomene* (French Guiana).

**Table S12.** List of ATAC-seq samples used in this study, and corresponding accession numbers

| Accession | Country | Genus | Species | Subsp |
|---|---|---|---|---|
| **ERS5932155** | Panama | Heliconius | erato | demophoon |
| **ERS5932158** | Panama | Heliconius | erato | hydara |
| **ERS5932159** | Panama | Heliconius | erato | hydara |
| **ERS5932160** | Panama | Heliconius | erato | hydara |
| **ERS5932156** | Panama | Heliconius | erato | demophoon |
| **ERS5932164** | Panama | Heliconius | melpomene | rosina |
| **ERS5932158** | Panama | Heliconius | melpomene | melpomene |
| **ERS5932165** | Panama | Heliconius | melpomene | rosina |
| **ERS5932162** | Panama | Heliconius | melpomene | melpomene |
| **ERS5932157** | Panama | Heliconius | erato | demophoon |
| **ERS5932163** | Panama | Heliconius | melpomene | melpomene |
| **ERS5932166** | Panama | Heliconius | melpomene | rosina |
