## Supplementary File 13 for "*Cortex cis*-regulatory switches establish scale colour identity and pattern diversity in *Heliconius*"

**Supplementary File 13:** Hi-C data shows looping of the CREs to the *cortex* promoter.

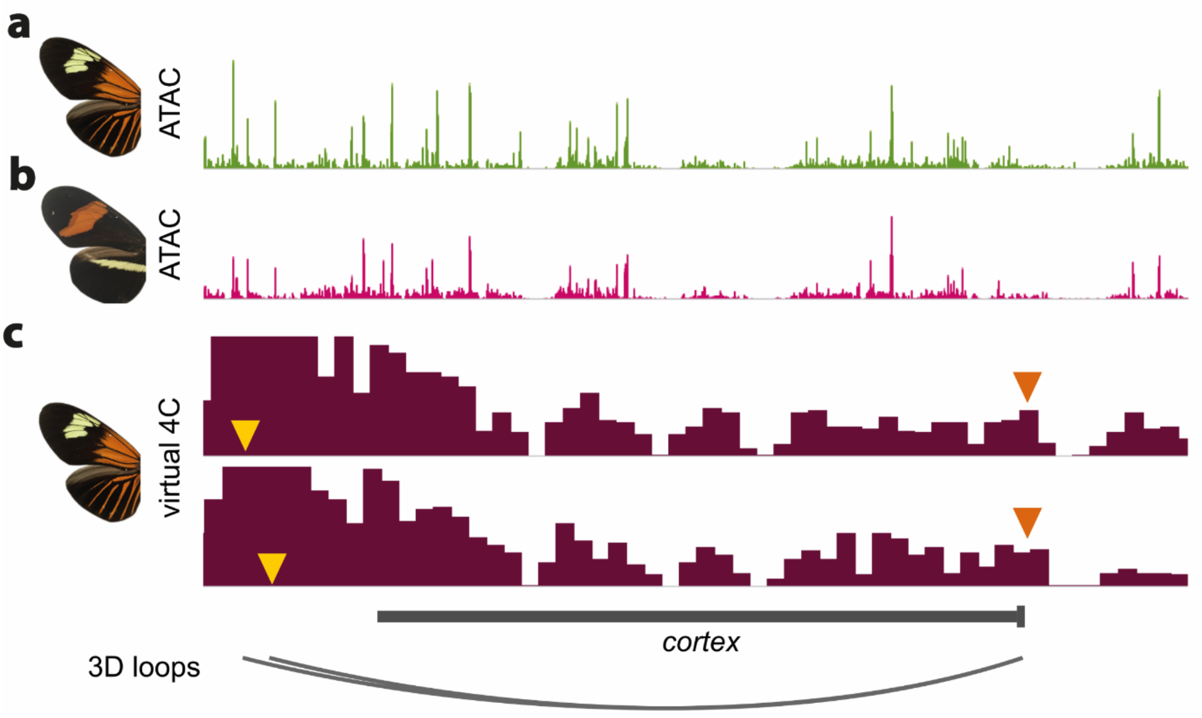


**Figure S13**. ATAC-Seq traces at the *cortex* locus for day 3 old pupal wings in *H. erato lativitta* **(a)** and *H. erato demophoon* **(b)**. Virtual 4C plots showing significant interactions ($p<0.05$) for the CREs assayed in the CRISPR experiments (yellow arrowheads) with the *cortex* promoter (orange arrowheads). Data from Lewis et al., 2019.
