## Supplementary File 14 for "*Cortex cis*-regulatory switches establish scale colour identity and pattern diversity in *Heliconius*"

**Supplementary File 14:** No evidence of deletion at the yellow bar CRE in *H. erato* populations.

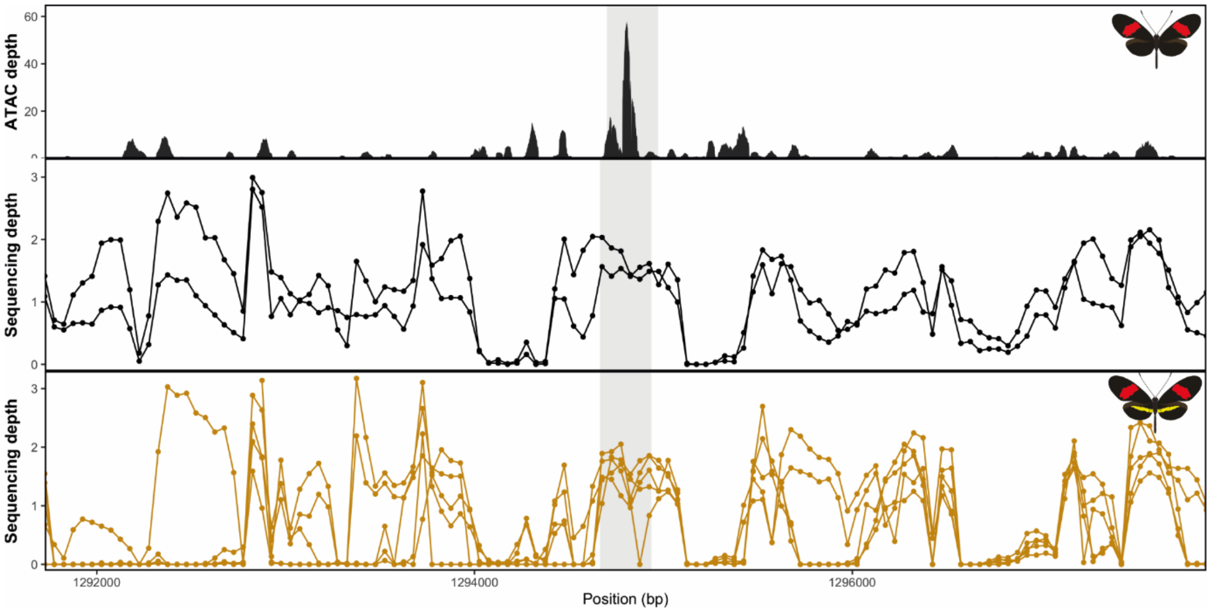


**Figure S14.1:** Mean sequence depth for *ATAC-Seq* in *H.erato* (top) and normalised sequencing depth for black hindwing populations (middle) and yellow barred populations (bottom). Sequences were mapped against the *H. erato lativitta* reference.

**Table S14.1.** List of individuals used in coverage depth analysis, and corresponding accession numbers

| Accession | Country | Genus | Species | Subsp |
|---|---|---|---|---|
| ERS4212889 | Colombia | Heliconius | melpomene | bellula |
| ERS4212890 | Colombia | Heliconius | melpomene | bellula |
| ERS4212589 | Colombia | Heliconius | melpomene | bellula |
| ERS4212891 | Colombia | Heliconius | melpomene | bellula |
| ERS4212892 | Colombia | Heliconius | melpomene | bellula |
| ERS2196350 | Colombia | Heliconius | melpomene | malleti |
| ERS1030547 | Colombia | Heliconius | melpomene | malleti |
| ERS1030548 | Colombia | Heliconius | melpomene | malleti |
| ERS2196351 | Colombia | Heliconius | melpomene | malleti |
| ERS1030546 | Colombia | Heliconius | melpomene | malleti |
| ERS4212641 | Colombia | Heliconius | timareta | tristero |
| ERS4212642 | Colombia | Heliconius | timareta | tristero |
| ERS4212643 | Colombia | Heliconius | timareta | tristero |
| ERS4212644 | Colombia | Heliconius | timareta | tristero |
| ERR2298254 | Peru | Heliconius | melpomene | amaryllis |
| ERR2298255 | Peru | Heliconius | melpomene | amaryllis |
| ERR2298256 | Peru | Heliconius | melpomene | amaryllis |
| ERR2298257 | Peru | Heliconius | melpomene | amaryllis |
| ERR2298260 | Peru | Heliconius | melpomene | amaryllis |
| ERR2298261 | Peru | Heliconius | melpomene | amaryllis |
| ERR2298205 | Panama | Heliconius | melpomene | rosina |
| ERR2298206 | Panama | Heliconius | melpomene | rosina |

| | | | | |
|---|---|---|---|---|
| ERR2298207 | Panama | Heliconius | melpomene | rosina |
| ERR2298208 | Panama | Heliconius | melpomene | rosina |
| ERR2298209 | Panama | Heliconius | melpomene | rosina |
| ERR2298210 | Panama | Heliconius | melpomene | rosina |
| ERR2298211 | Panama | Heliconius | melpomene | rosina |
| ERR2298219 | Panama | Heliconius | melpomene | rosina |
| ERR2298221 | Panama | Heliconius | melpomene | rosina |
| ERR2298262 | Peru | Heliconius | timareta | thelxinoe |
| ERR2298263 | Peru | Heliconius | timareta | thelxinoe |
| ERR2298264 | Peru | Heliconius | timareta | thelxinoe |
| ERR2298265 | Peru | Heliconius | timareta | thelxinoe |
| ERR2298266 | Peru | Heliconius | timareta | thelxinoe |
| ERS4212893 | Colombia | Heliconius | erato | dignus |
| ERS4212895 | Colombia | Heliconius | erato | dignus |
| ERS4212896 | Colombia | Heliconius | erato | dignus |
| ERS4212581 | Colombia | Heliconius | erato | dignus |
| ERS4212897 | Colombia | Heliconius | erato | dignus |
| SRR4031993 | Colombia | Heliconius | erato | chestertonii |
| SRR4032020 | Colombia | Heliconius | erato | chestertonii |
| SRR4032031 | Colombia | Heliconius | erato | chestertonii |
| SRR4032104 | Colombia | Heliconius | erato | chestertonii |
| SRR4032105 | Colombia | Heliconius | erato | chestertonii |
| SRR4031996 | Colombia | Heliconius | erato | hydara |
| SRR4031999 | Colombia | Heliconius | erato | hydara |
| SRR4032000 | Colombia | Heliconius | erato | hydara |
| SRR4032061 | Colombia | Heliconius | erato | hydara |
| SRR4032068 | Colombia | Heliconius | erato | hydara |
| SRR4031995 | Panama | Heliconius | erato | demophoon |
| SRR4031997 | Panama | Heliconius | erato | demophoon |
| SRR4032001 | Panama | Heliconius | erato | demophoon |
| SRR4032002 | Panama | Heliconius | erato | demophoon |
| SRR4032093 | Panama | Heliconius | erato | demophoon |
| SRR4032044 | Ecuador | Heliconius | erato | lativitta |
| SRR4032045 | Ecuador | Heliconius | erato | lativitta |
| SRR4032046 | Ecuador | Heliconius | erato | lativitta |
| SRR4032047 | Ecuador | Heliconius | erato | lativitta |
| SRR4032053 | Ecuador | Heliconius | erato | lativitta |
| SRR4032032 | Peru | Heliconius | erato | favorinus |
| SRR4032056 | Peru | Heliconius | erato | favorinus |
| SRR4032057 | Peru | Heliconius | erato | favorinus |
| SRR4032058 | Peru | Heliconius | erato | favorinus |
| SRR4032059 | Peru | Heliconius | erato | favorinus |
| SRR4032003 | Brazil | Heliconius | erato | phyllis |
| SRR4032004 | Brazil | Heliconius | erato | phyllis |
| SRR4032006 | Brazil | Heliconius | erato | phyllis |
| SRR4032007 | Brazil | Heliconius | erato | phyllis |
| ERR2298201 | French Guiana | Heliconius | melpomene | melpomene |
| ERR2298204 | French Guiana | Heliconius | melpomene | melpomene |

| | | | | |
|---|---|---|---|---|
| ERR2298207 | French Guiana | Heliconius | melpomene | melpomene |
| ERR2298214 | French Guiana | Heliconius | melpomene | melpomene |
| ERR2298216 | French Guiana | Heliconius | melpomene | melpomene |
| ERR2298235 | Colombia | Heliconius | timareta | florencia |
| ERR2298233 | Colombia | Heliconius | timareta | florencia |
| ERR2298234 | Colombia | Heliconius | timareta | florencia |
| ERR2298237 | Colombia | Heliconius | timareta | florencia |
| ERR2298236 | Colombia | Heliconius | timareta | florencia |

**Supplementary Text 14.1**

Consensus sequences recovered from Sanger sequencing across the *H. melpomene/timareta* CRE. The BovB-like TE element is indicated in blue; The Helitron-like fragment in orange. Both are absent from the *H. melpomene melpomene* sequence.

>*H. melpomene rosina* CRE

ACTTCCAACATTCCGCAACATTTCATATTTGGATAGACCACCTTCGCTAATCAGTTATCAGTAATTA
AAATGTACATAATTGTATGAACAAACAGCTTACAATGTAAGGAGTGGAAAACAATATAAGTAAAA
CAATATAAGTAAAGTTTCCTCGTATAAATTCATATAACGTTACGTTCAGGTTATTATTTATTTTAAG
TACAAATGATGATAAGACTAATGATATGCTGTAATATGTTGCATATAGGGTATCTTCCTCCTGGCA
TTAATCCCGGCTATTGCCAGGGTCTGCCCTCCTACTCAACCTACTCCACTTTGCACGGTCTTTTGCG
TCCTCGGTGGTTAGACCATTGGCTCTCATGTCCTGCATCACGACATCCAGCCAGCGCTTCCTTGGGC
GTCGAGGAGGTCTCTCTCCAGGAACTGATATAGAAAGGCACTTGCATCCGACATAGTTCTTAGGCC
GGCGCGTACCATGGCCAAACCATTTCAGACGACGCTCTTGGAGCTTATCACGGACACGGACGGAC
CCCAAGACTACCTCGAATGAATGTGTTGCGTATGCGGTCAGACCGCGTTACGCCGCACATCCACCT
CAGCATCTTTATTTCCGTGACGTGAAGCTCCTGAGTGTGCCGAGATAGTGCCGGCCAACATTCGCT
GCCGTATAAAAGAACCGGTCGGATGATGCTCTTGTATATCAGCCCCTTGAGCTTGGGCGGTATTCT
GCGGTCGCAGACCACACCAGTGACCTCCCGCCATTTGGCCCAGGCAGCGCTTATCCGGCCTTGGAC
ATCGTGATCGATGCCTCCAGACTCGTGCATAACGGTTCCAAGGTACCTGAACTTTTCCGACTTAAC
GGCTGGCTCAGGACCTATAAGGATCGTGCTCGAGTCCGGGCTCCCGCAGGCTATGGCCATGGCCAT
GTTGCATATAGGGTATATTTTTCAATACTAAGGATTTTGGTGGTCTATCAATTAAAATAAAATTTTC
TATGTTAATTATCTTACTGTTATATTTTTTCGATTTTATACCTAACTAGCGACCCTCTTGCGGCTTCG
CCCGCTTTTACTACTGGATTATTCATATAATGTATGCTTGCAAAGCACTTAAGATAATGTGAAAATT
ATTTAAACCCTAATGCAACCCGCATTTTCGTAGTTACTGCTACTTCATTATATTCATTTTTTAACTTT
TAATGAACGTTATCATTAAGTCTTATGGCAATTTTGTATTGAAACAAAACTGATCGTAATTTTTTCT
AAAAAAAAATACACAGAAATCATTTTATTAAAAAGATATAAAGCCAAATAATAAATAAAAGTTGT
ATAATACAGTATAACGTACATAGAAAATTTAAATAACCTGAGTTACACCACTTTGCCTAGACGCCC
CGCACTGGGTGGCTGACGTCAATT

>*H. melpomene amaryllis* CRE

ACTTCGCTATCAGTTATCAGTAATTAAAATGTACATAATTGTATGAACAAACAGCTTACAATGTAA
GGAGTGGAAAACAATATAAGTAAAACAATATAAGTAAAGTTTCCTCGTATAAATTCATATAACGT
TACGTTCAGGTTATTATTTATTTTAAGTACAAATGATGATAAGACTAATGATATGCTGTAATATGTT
GCATATAGGGTATCTTCCTCCTGGCATTAATCCCGGCTATTGCCAGGGTCTGCCCTCCTACTCAACC
TACTCCACTTTGCACGGTCTTTTGCGTCCTCGGTGGTTAGACCATTGGCTCTCATGTCCTGCATCAC
GACATCCAGCCAGCGCTTCCTTGGGCGTCGAGGAGGTCTCTCTCCAGGAACTGATATAGAAAGGC
ACTTGCATCCGACATAGTTCTTAGGCCGGCGCGTACCATGGCCAAACCATTTCAGACGACGCTCTT
GGAGCTTATCCGCTACGTCACGGACCCCAAGACTACCTCGAATGAATGTGTTGCGTATGCGGTCAG
ACCGCGTTACGCCGCMCATCCACCTCAGCATCTTTATTTCCGTGACGTGAAGCTCCTGAGTGTGCC
GAGATAGTGCCGGCCAACATTCGCTGCCGTATAAAAGAACCGGTCGGATRATGCTCTTGTATATCA
GCCCCTTGAGCTTGGGCGGTATTCTGCGGTCGCAGACCACACCAGTGACCTCCCGCCATTTGGCCC
AGGCAGCGCTTATCCGGCCTTGGACATCGTGATCGATGCCTCCAGACTCGTGCATAACGGTTCCAA
GGTACCTGAACTTTTCCGACTTAACGGCTGGCTCAGGACCTATAAGGATCGTGCTCGAGTCCGGGC

TCCCGCAGGCCATGGCCATGGCCATGTTGCATATAGGGTATATTTTTCAATACTAAGGATTTTGGT
GGTCTATCGATTAAAATAAAATTTTCTATGTTAATTATCTTACTGTTATATTTTTTCGATTTTATACC
TAACTAGCGACCCTCTTGCGGCTTCGCCCGCTTTTACTACTGGATTATTTATATAATGTATGCTTGC
AAAGCATTTAACATAATGTGAAAATTATTTAAACCCTAATGCAACCCGCATTTTCGTAGTTACTGC
TACTTCATTATATTCATTTTTTAACTTTTAATGAACGTTATCATTAAGTCTTATGGWAATTTTGTATT
GAAACAAAACTGATCGTAATTTTTTCTAAAAAAAAATACACAGAAATCATTTTATTAAAAAGATAT
AAAGCCAAATAATAAATAAAAGTTGTATAATACAGTATAACGTACATAGAAAATTTAAATAACCT
GAGTACACCA

>*H. melpomene bellula* CRE

CACTTCGCTATCAGTTATCAGTAATTAAAATGTACATAATTGTATGAACAAACAGCTTACAATGTA
AGGAGTGGAAAACAATATAAGTAAAACAATATAAGTAAAGTTTCCTCGTATAAATTCATATAACG
TTACGTTCAGGTTATTATTTATTTTAAGTACAAATGATGATAAGACTAATGATATGCTGTAATATGT
TGCATATAGGGTATCTTCCTCCTGGCATTAATCCCGGCTATTGCCAGGGTCTGCCCTCCTACTCAAC
CTACTCCACTTTGCACGGTCTTTTGCGTCCTCGGTGGTTAGACCATTGGCTCTCATGTCCTGCATCA
CGACATCCAGCCAGCGCTTCCTTGGGCGTCGAGGAGGTCTCTCTCCAGGAACTGATATAGAAAGG
CACTTGCATCCGACATAGTTCTTAGGCCGGCGCGTACCATGGCCAAACCATTTCAGACGACGCTCT
TGGAGCTTATCCGCTACGTCACGGACCCCAAGACTACCTCGAATGAATGTGTTGCGTATGCGGTCA
GACCGCGTTACGCCGCCCATCCACCTCAGCATCTTTATTTCCGTGACGTGAAGCTCCTGAGTGTGC
CGAGATAGTGCCGGCCAACATTCGCTGCCGTATAAAAGAACCGGTCGGATAATGCTCTTGTATATC
AGCCCCTTGAGCTTGGGCGGTATTCTGCGGTCGCAGACCACACCAGTGACCTCCCGCCATTTGGCC
CAGGCAGCGCTTATCCGGCCTTGGACATCGTGATCGATGCCTCCAGACTCGTGCATAACGGTTCCA
AGGTACCTGAACTTTTCCGACTTAACGGCTGGCTCAGGACCTATAAGGATCGTGCTCGAGTCCGGG
CTCCCGCAGGCCATGGCCATGGCCATGTTGCATATAGGGTATATTTTTCAATACTAAGGATTTTGG
TGGTCTATCGATTAAAATAAAATTTTCTATGTTAATTATCTTACTGTTATATTTTTTCGATTTTATAC
CTAACTAGCGACCCTCTTGCGGCTTCGCCCGCTTTTACTACTGGATTATTTATATAATGTATGCTTG
CAAAGCATTTAACATAATGTGAAAATTATTTAAACCCTAATGCAACCCGCATTTTCGTAGTTACTG
CTACTTCATTATATTCATTTTTTAACTTTTAATGAACGTTATCATTAAGTCTTATGGTAATTTTGTAT
TGAAACAAAACTGATCGTAATTTTTTCTAAAAAAAAATACACAGAAATCATTTTATTAAAAAGATA
TAAAGCCAAATAATAAATAAAAGTTGTATAATACAGTATAACGTACATAGAAAATTTAAATAACC
TGAGTACACCA

>*H. timareta tristero* CRE

TTCGCTATCAGTTATCAGTAATTAAAATGTACATAATTGTATGAACAAACAGCTTACAATGTAAGG
AGTGGAAAACAATATAAGTAAAACAATATAAGTAAAGTTTMCTCGTATAAATTCATATAACGTTA
CGTTCAGGTTATTATTTATTTTAAGTACAAATGATGATAAGACTAATGATATGCTGTAAT

ATGTTGCATATAGGGTATCTTCCTCCTGGCATTAATCCCGGCTATTGCCAGGGTCTGCCCTCCTACT
CAACCTACTCCACTTTGCACGGTCTTTTGCGTCCTCGGTGGTTAGACCATTGGCTCTCATGTCCTGC
ATCACGACATCCAGCCAGCGCTTCCTTGGGCGTCGAGGAGGTCTCTCTCCAGGAACTGATATAGAA
AGGCACTTGCATCCGACATAGTTCTTAGGCCGGCGCGTACCATGGCCAAACCATTTCAGACGACGC
TCTTGGAGCTTATCCGCTACGTCACGGACCCCAAGACTACCTCGAATGAATGTGTTGCGTATGCGG
TCAGACCGCGTTACGCCGCACATCCACCTCAGCATCTTTATTTSMGTGACGTGAAGCTCCTGAGTG
TGCCGAGATAGTGCCGGCCAACATTCGCTGCCGTATAAAAGAACCGGTCGGATGATGCTCTTGTAT
ATCAGCCCCTTGAGCTTGGGCGGTATTCTGCGGTCGCAGACCACACCRGTGACCTCCCGCCATTTG
GCCCAGGCAGCGCTTATCTGGCCTTGGACATCGTGATCGATGCCTCCAGACTCGTGCATAACGGTT
CCAAGGTACCTGAACTTTTCCGACTTAACGGCTGGCTCAGGACCTATAAGGATCGTGCTCGAGTCC
GGGCTCCCGCAGGYYATGGCCATGGCCATGTTGCATATAGGGTATATTTTTCAATACTAAGGATTT
TGGTGGTCTATCARTTAAAATAAAATGTTCTATGTTAATTATCTTACTGYTATATTTTTTCGAYYYT
ATACCTADCTAGCGAYCCTCWTGCRRCTTCGSCYGNCTAAGCACTTAAGATAATGTGAAAATTATT
TAAACCCTAATGCAACCCGCATTTTYGTAGTTACTGCTACTTCATTATATTCATTTTTTAACTTTTAA
TGAACGTTATCATTAAGTCTTATGGMAATTTTGTATTGAAACAAAACTGATCGTAATTTTTTYTAA
AAAAAAATACACAGAAATCATTTTATTAAAAAGATATAAAGCCAAATAATAAATAAAAGTTGTAT
AATACAGTATAACGTACATAGAAAATTTAAATAACCTGAGTAC

>*H. melpomene melpomene* CRE

ACTTCCAACATTCCGCAACATTTCTTATTTGGTTAGACATTAGACCAACATCGCTAATCATTTATCA
ATAATTAAAATGTACATAATTGTATGAACAAACAGCTTACAATATAAGGATGGAACTAAATAAAG
TTTTATTATTATTATTTTTTATTTATGGAAAACAATTCATACATTATTAAGAGTAATAAATAAGTAA
AGTTTCCTCGTATAAATTCATATAACGTTACGTTCAAGTTATTATTTATTTTAAGTACAAATGATGA
TAACACTAAAAATATGCTGTAATATGTTGCATATAGGGTATATTTTTCAATACTAAGGAAAATGGT
GGTCTATCAATTAAAATAAAATTTTCTATGTTAATTATCTTACACACCTTACATCTAGTTATATTTTT
TCGATTGTATACCTAATCAACCATTAATTAGTATTCAAAAGTATGATAATTTCTACATAACGCGAA
AAAAGGTACCGTTTTCAAGAAGCATGTGTTCTAGTCCTTTTCTTTCCTTAGAGTATGAATAATACAT
TAAATAAAGGTGGCATGGCATGGTTCATGGCTGCTACTTCATTATATTCATTTATTAACTTTTATGA
CGTTCATTAAGTCTTATGGCAATTTTGTATTGAAACAAAACTTATTGTAATTTTTTCTAAAAAAAAT
ACACAGGAATCGTTTTATTAAAAAGATATAAAGCCAAATAATAAATAAAACTTGTATAATACAGT
ATAACGTACGTGGAAAATTTAAATAACCTGAGTTACACCACTTTGCTTGGACGCCCCGCACTGGGT
GGCTGACGTCAATT

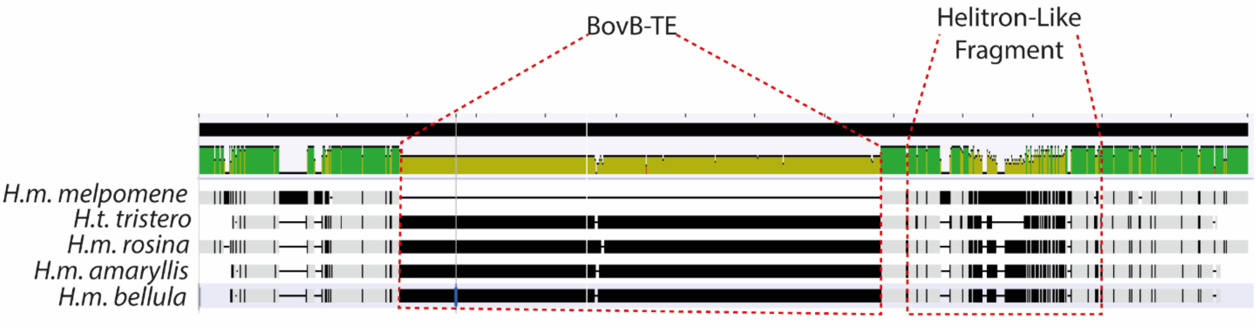


**Figure S14.2:** Alignment visualisation of the sequences above. The BovB-TE insertion is evident in all yellow-barred morphs, as well as the divergent Helitron like sequence. This is absent from the black hindwing morph (top, *H.m. melpomene*).
