## Supplementary File 15 for "*Cortex cis*-regulatory switches establish scale colour identity and pattern diversity in *Heliconius*"

### Supplementary File 15: Quantative measures of scale structures and features

### Table S15a: Pairwise Wilcox test adjusted P values for quantative measures of scale structures and features in *H. melpomene*

**MELPOMENE SCALE WIDTH**

| | cythera, black WT | cythera, white WT | cythera, white Mutant | rosina, black WT | rosina, yellow mutant |
|---|---|---|---|---|---|
| **cythera, white WT** | 0.25579 | - | - | - | - |
| **cythera, white Mutant** | 0.40818 | 0.60473 | - | - | - |
| **rosina, black WT** | 0.00063 | 0.03835 | 0.00047 | - | - |
| **rosina, yellow mutant** | 0.00936 | 0.40317 | 0.03835 | 0.00047 | - |
| **rosina, yellow WT** | 0.00067 | 0.07611 | 0.01998 | 0.10855 | 0.25579 |

**MELPOMENE WIDTH**

**MELPOMENE SCALE LENGTH**

| | cythera, black WT | cythera, white WT | cythera, white Mutant | rosina, black WT | rosina, yellow mutant |
|---|---|---|---|---|---|
| **cythera, white WT** | 0.00253 | - | - | - | - |
| **cythera, white Mutant** | 0.419 | 0.04351 | - | - | - |
| **rosina, black WT** | 1.90E-06 | 3.00E-06 | 0.00012 | - | - |
| **rosina, yellow mutant** | 5.50E-08 | 7.80E-08 | 2.10E-05 | 0.01305 | - |
| **rosina, yellow WT** | 3.40E-05 | 9.30E-06 | 0.0003 | 0.02436 | 0.91489 |

**MELPOMENE SERRATIONS**

| | cythera, black WT | cythera, white WT | cythera, white Mutant | rosina, black WT | rosina, yellow mutant |
|---|---|---|---|---|---|
| **cythera, white WT** | 0.4766 | - | - | - | - |
| **cythera, white Mutant** | 1 | 0.4489 | - | - | - |
| **rosina, black WT** | 0.1147 | 0.674 | 0.0575 | - | - |
| **rosina, yellow mutant** | 0.6758 | 0.6758 | 0.5145 | 0.4164 | - |
| **rosina, yellow WT** | 0.0084 | 0.0815 | 0.0036 | 0.0867 | 0.0120 |

**MELPOMENE RIDGE PERIODICITY**

| | cythera, black WT | cythera, white WT | cythera, white Mutant | rosina, black WT | rosina, yellow mutant |
|---|---|---|---|---|---|
| **cythera, white WT** | 0.05492 | - | - | - | - |
| **cythera, white Mutant** | 0.92072 | 0.11904 | - | - | - |
| **rosina, black WT** | 0.18957 | 0.01475 | 0.78471 | - | - |
| **rosina, yellow mutant** | 0.00819 | 0.00175 | 0.11904 | 0.044 | - |
| **rosina, yellow WT** | 0.00018 | 0.00029 | 0.0056 | 0.00018 | 0.01524 |

**MELPOMENE CROSSRIB PERIODICITY**

| | cythera, black WT |
|---|---|
| **rosina, black WT** | 7.1e-05 |

**MELPOMENE MICRORIB PERIODICITY**

| | cythera, white WT | cythera, white Mutant | rosina, yellow mutant |
|---|---|---|---|
| **cythera, white Mutant** | 0.083 | - | - |
| **rosina, yellow mutant** | 0.667 | 0.083 | - |
| **rosina, yellow WT** | 0.083 | 0.686 | 0.083 |

### Supplementary File 15: Quantative measures of scale structures and features

### Table S15b: Pairwise Wilcox test adjusted P values for quantative measures of scale structures and features in *H. erato*

**ERATO SCALE WIDTH**

| | cyrbia, black WT | cyrbia, white WT | cyrbia, white Mutant | demophoon, black WT | demophoon, yellow Mutant | demophoon, yellow WT |
|---|---|---|---|---|---|---|
| **cyrbia, white WT** | 6.00E-05 | - | - | - | - | - |
| **cyrbia, white Mutant** | 0.52849 | 0.00057 | - | - | - | - |
| **demophoon, black WT** | 0.51944 | 0.00015 | 0.37403 | - | - | - |
| **demophoon, yellow Mutant** | 0.60363 | 6.70E-05 | 0.50041 | 0.51944 | - | - |
| **demophoon, yellow WT** | 0.27121 | 2.00E-06 | 0.86485 | 0.14757 | 0.09625 | - |
| **hydara, red WT** | 0.01967 | 0.00057 | 0.85909 | 0.0102 | 0.04998 | 0.85909 |

**ERATO SCALE LENGTH**

| | cyrbia, black WT | cyrbia, white WT | cyrbia, white Mutant | demophoon, black WT | demophoon, yellow Mutant | demophoon, yellow WT |
|---|---|---|---|---|---|---|
| **cyrbia, white WT** | 0.3587 | - | - | - | - | - |
| **cyrbia, white Mutant** | 0.4991 | 0.8479 | - | - | - | - |
| **demophoon, black WT** | 0.0014 | 0.4521 | 0.3587 | - | - | - |
| **demophoon, yellow Mutant** | 0.4958 | 0.0961 | 0.1443 | 0.0002 | - | - |
| **demophoon, yellow WT** | 0.093 | 0.0206 | 0.1021 | 1.50E-05 | 0.4397 | - |
| **hydara, red WT** | 0.1443 | 0.8479 | 1 | 0.1306 | 0.075 | 0.0176 |

**ERATO PRONG NUMBER**

| | cyrbia, black WT | cyrbia, white WT | cyrbia, white Mutant | demophoon, black WT | demophoon, yellow Mutant | demophoon, yellow WT |
|---|---|---|---|---|---|---|
| **cyrbia, white WT** | 0.00028 | - | - | - | - | - |
| **cyrbia, white Mutant** | 0.15704 | 0.00241 | - | - | - | - |
| **demophoon, black WT** | 0.00151 | 0.00014 | 0.04219 | - | - | - |
| **demophoon, yellow Mutant** | 0.03611 | 6.00E-05 | 0.12531 | 0.06224 | - | - |
| **demophoon, yellow WT** | 0.06115 | 6.00E-05 | 0.80401 | 0.00655 | 0.06115 | - |
| **hydara, red WT** | 0.01286 | 0.00241 | 0.58257 | 0.06224 | 0.19112 | 0.69988 |

**ERATO RIDGE PERIODICITY**

| | cyrbia, black WT | cyrbia, white WT | cyrbia, white Mutant | demophoon, black WT | demophoon, yellow Mutant | demophoon, yellow WT |
|---|---|---|---|---|---|---|
| **cyrbia, white WT** | 0.02624 | - | - | - | - | - |
| **cyrbia, white Mutant** | 0.00728 | 0.05495 | - | - | - | - |
| **demophoon, black WT** | 0.3655 | 0.01305 | 0.00036 | - | - | - |
| **demophoon, yellow Mutant** | 0.00189 | 0.01142 | 0.50312 | 0.00094 | - | - |
| **demophoon, yellow WT** | 0.25182 | 0.00582 | 0.00021 | 0.06393 | 0.00021 | - |
| **hydara, red WT** | 0.02008 | 0.05495 | 0.05495 | 0.06247 | 0.10949 | 0.00617 |

**ERATO CROSSRIB PERIODICITY**

| | cyrbia, black WT | demophoon, black WT |
|---|---|---|
| **demophoon, black WT** | 0.00042 | - |
| **hydara, red WT** | 0.00163 | 0.00015 |

**ERATO MICRORIB PERIODICITY**

| | wt_white.cyr | mut_white.cyr | mut_yellow.dem |
|---|---|---|---|
| **mut_white.cyr** | 0.194 | - | - |
| **mut_yellow.dem** | 0.398 | 0.801 | - |
| **wt_yellow.dem** | 0.013 | 0.194 | 0.27 |

**Supplementary File 15: Quantative measures of scale structures and features**

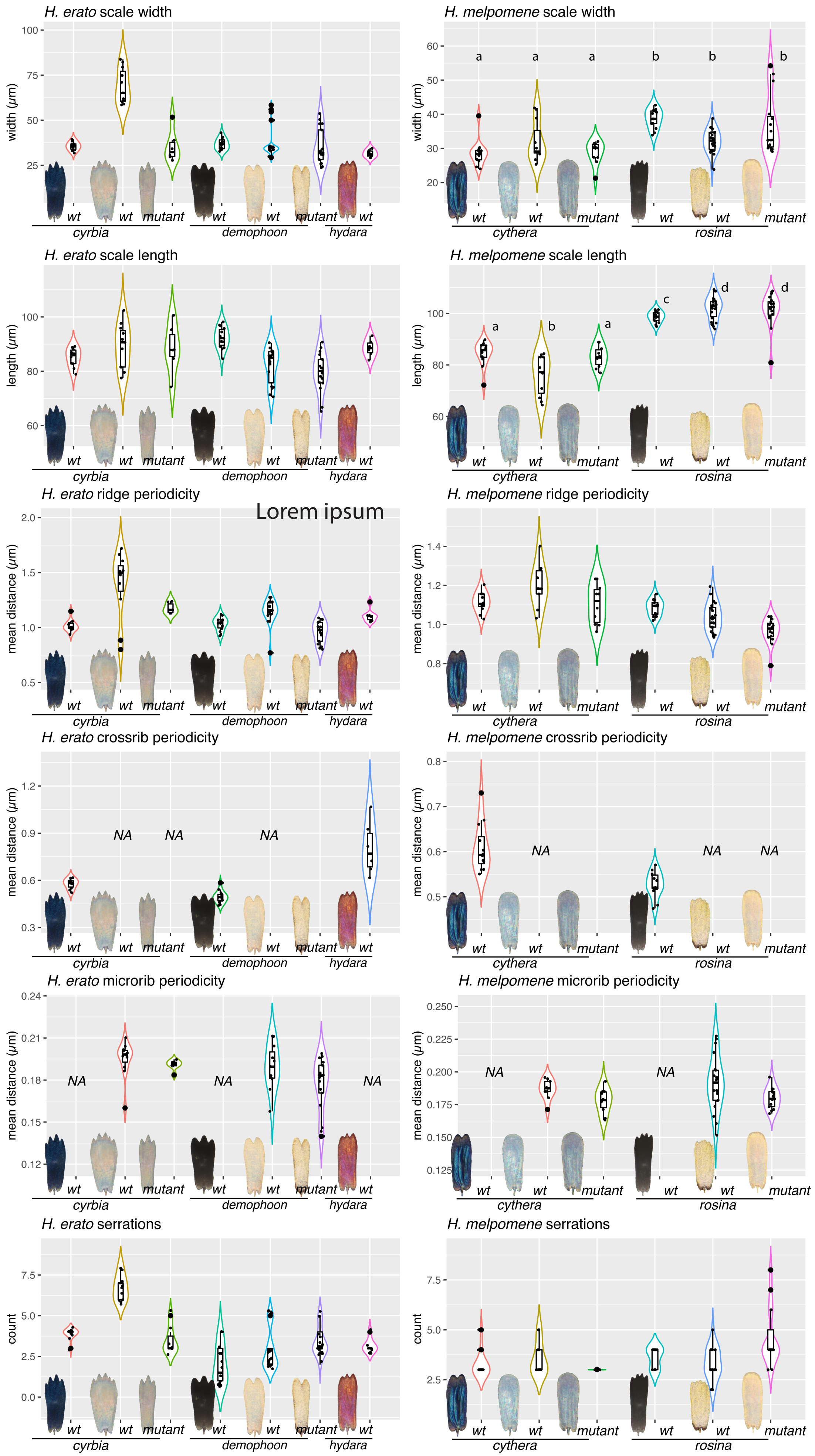

**Supplementary File 15: Quantative measures of scale structures and features**

**Figure S15: Quantitative measures of scale structures and features**.

For each scale type depicted in figure 9, between 8 and 15 scales were removed from the wing with an eyelash tool and imaged on SEM at 3000x. (Raw EM stitches can be found at the Dryad entry for this manuscript). Each measurement was taken as previously described by Day et al (2019). Briefly, a line segment was drawn over the scale in FIJI, and pixel intensity levels extracted. The resulting curves were subject to Fourier analysis to get mean measures for Ridge periodicity, crossrib periodicity and microrib periodicity. Serrations were counted manually.
